# *Clostridioides difficile* major toxins remodel the intestinal epithelia, affecting spore adherence/internalization into intestinal tissue and their association with gut vitronectin

**DOI:** 10.1101/2025.01.29.635439

**Authors:** Pablo Castro-Cordova, Osiris K. Lopez-Garcia, Josué Orozco, Nicolás Montes-Bravo, Fernando Gil, Marjorie Pizarro-Guajardo, Daniel Paredes-Sabja

**Affiliations:** Millennium Nucleus in the Biology of Intestinal Microbiota, Santiago, Chile; IMPACT, Center of Interventional Medicine for Precision and Advanced Cellular Therapy, Santiago, Chile; Laboratory of Nano-Regenerative Medicine, Centro de Investigación e Innovación Biomédica (CiiB), Faculty of Medicine, Universidad de los Andes, Chile; Interdisciplinary Program in Genetics & Genomics, Texas A&M University, College Station, TX USA; Department of Biology, Texas A&M University, College Station, TX USA; Microbiota-Host Interactions & Clostridia Research Group, Universidad Andres Bello, Santiago, Chile; Department of Biology, Texas A&M University, College Station, TX USA; dparedes-

## Abstract

The most common cause of healthcare-associated diarrhea and colitis in the U.S., is *Clostridioides difficile*, a spore-forming pathogen. Two toxins, TcdA and TcdB, are major virulence factors essential for disease manifestations, while *C. difficile* spores are essential for disease transmission and recurrence. Both toxins cause major damage to the epithelial barrier, trigger massive inflammation, and reshape the microbiome and metabolic composition, facilitating *C. difficile* colonization. *C. difficile* spores, essential for transmission and recurrence of the disease, persist adhered and internalized in the intestinal epithelia. Studies have suggested that toxin-neutralization in combination with antibiotic during CDI treatment in humans significantly reduces disease recurrence, suggesting a link between toxin-mediated damage and spore persistence. Here, we show that TcdA/TcdB-intoxication of intestinal epithelial Caco-2 cells leads to remodeling of accessible levels of fibronectin (Fn) and vitronectin (Vn) and their cognate alpha-integrin subunits. While TcdB-intoxication of intestinal tissue had no impact in accessible levels of Fn and Vn, but significantly increased levels of intracellular Vn. We observed that Fn and Vn released to the supernatant readily bind to *C. difficile* spores *in vitro*, while TcdB-intoxication of intestinal tissue led to increased association of *C. difficile* spores with gut Vn. Toxin-intoxication of the intestinal tissue also contributes to increased adherence and internalization of *C. difficile* spores. However, TcdB-intoxicated ligated loops infected of mice treated with Bezlotoxumanb (monoclonal anti-TcdB antibodies) did not prevent TcdB-mediated increased spore adherence and internalization into intestinal tissue. This study highlights the importance of studying the impact of *C. difficile* toxins of host tissues has in *C. difficile* interaction with host surfaces that may contribute to increased persistence and disease recurrence.

## Introduction

Frequently related to antibiotic-associated diarrhea (AAD) is *Clostridioides difficile*, which accounts for approximately 30% of cases^1^. *C. difficile* infections (CDI) have emerged as a significant concern for healthcare systems globally, occasionally leading to severe outbreaks with mortality rates as high as 20%. Most patients typically respond favorably to medical treatments like vancomycin and metronidazole^1, 2^. However, in contrast to other gastrointestinal infections, CDI exhibits an exceptionally high recurrence rate, ranging from 20% after the first episode, to 40% after the second, and even 60% after the third, potentially leading to the development of more severe symptoms^1, 3, 4^. In Europe and the USA alone, CDI results in substantial economic losses estimated at around US$ 5 billion annually for the healthcare system^1, 5^. Addressing recurrent CDI presents a considerable clinical challenge.

In response to a dysbiotic microbiota (commonly caused by antibiotics), ingested or indigenous spores will germinate leading to *C. difficile* colonization and infection^6^. Two main virulence factors are responsible for the clinical manifestations of the disease, toxins TcdA and TcdB^7, 8^. Both toxins monoglucosylate host GTPases of the Rho family resulting in downstream cellular changes as tight junction collapse, detachment of cells, and impairment of the intestinal epithelial barrier^7, 8^. In addition, exposure to toxins triggers intestinal epithelial cells to release several pro-inflammatory cytokines, resulting in attraction of neutrophils and acute mucosal inflammation^7, 8^. These events culminate in further activation of immune cells, exacerbating inflammation and mucosal damage. In addition to these events, *C. difficile* initiates a sporulation cycle during infection, resulting in a metabolically dormant spore that is impervious to all known antibiotics and regarded as essential for recurrence of the infection ^9^.

Concatenated to the loose and tight mucosal layers in the gastrointestinal tract, is an array of potential interaction sites commonly exploited by enteric pathogens ^10^. Of these, extracellular matrix (ECM) proteins are widely exploited as binding molecular bridges by pathogens, specifically fibronectin (Fn) and vitronectin (Vn)^11, 12^. Fn, is a 450 kDa dimeric, multi-modular glycoprotein which can exist in two different forms, plasma and cellular Fn. Plasma Fn is synthetized by hepatocytes and secreted to the blood plasma to levels of 300 - 400 μg /ml ^13^. By contrast, cellular Fn is synthetized by various cell types, including fibroblasts, intestinal epithelial cells, endothelial cells, chondrocytes, synovial cells and myocytes ^14, 15^, and in the intestinal epithelial crypt cells ^13^. Vn is a multidomain glycoprotein of 75 kDa mainly synthetized in the liver and secreted in the plasma, and to a lesser extent produced in platelets and macrophages ^16^. Vn is highly abundant in the plasma (0.2 - 0.4 mg/ml) and in the extracellular matrix^16^, it interacts with several other molecules to regulate various functions including wound healing, the complement system, cell migration, and adhesion^17–19^.

Both, Fn and Vn are implicated in tissue repair. Inflammatory stimuli and intestinal injury derived from chemical-colitis leads to increase expression of intestinal Fn in the mucosal layers ^20, 21^, primarily by intestinal epithelial cells ^22^. Notably, patients with inflammatory bowel disease also exhibit increased Fn levels ^23^. Although, in experimental-colitis expression of Vn increases and is associated with protection against colitis ^24^. Evidence of Fn-remodeling by *C. difficile* binary toxin has been documented ^25, 26^; however, it is unclear how both *C. difficile* major toxins impact levels of Fn and Vn in intestinal epithelial cells.

*C. difficile* toxins, TcdA and TcdB, have traditionally been considered primary virulence factors essential for clinical manifestation of the disease^27^. However, the most recently FDA approved intravenous CDI therapeutic, Zinplava, which neutralizes *C. difficile* TcdB, was found to reduce the rate of recurrence of CDI^28^. Zinplava is an intravenously administered anti-TcdB monoclonal antibody (Bezlotoxumab) intended to be used in combination with antibiotics ^29–32^. Bezlotoxumab clinical trials results demonstrate that administration of anti-toxin antibodies correlates with lower rates of CDI recurrence ^29–32^, suggesting that toxin-mediated damage could be contributing indirectly to *C. difficile* persistence. Although TcdA/TcdB-intoxication of intestinal epithelial cells results in increased adherence and internalization of *C. difficile* spores *in vitro* ^33^, and increased levels of Fn in the colon during infection ^34, 35^, the underlying mechanisms remain unknown. Consequently, in this work we assessed how TcdA/TcdB-intoxication remodels Fn and Vn, and spore-epithelial cell interactions *in vivo*, and whether Bezlotoxumab can prevent toxin-mediated remodeling and spore-binding in the intestinal mucosa. Results demonstrate that while TcdA/TcdB-intoxication of intestinal epithelial cells *in vitro* increased levels of accessible Fn and Vn, this was not the case in an ileal loop murine model. Moreover, confocal micrographs show that while a slight increase in Fn- and Vn-spore interactions were observed at the mucosal layer, these were a small proportion of analyzed spores. Interestingly, TcdB-mediated intoxication led to increased adherence and internalization of *C. difficile* spores to the intestinal epithelial layer *in vivo*, interactions that were reduced by administration of Bezlotoxumab.

## Results

### TcdA/TcdB-intoxication increases accessible Fn and Vn in intestinal epithelial cells

In prior work, we demonstrated that Fn and Vn are required for spore-entry into intestinal epithelial cells (IECs) ^36^. TcdA/TcdB-intoxication of epithelial cells leads to opening of adherent junctions and increased spore internalization ^33^. Moreover, since intestinal inflammation and epithelial damage leads to increased intestinal Fn and Vn ^20, 21^, we explored how TcdA/TcdB-intoxication impacted redistribution of both molecules. For this, we quantified accessible and total levels of protein in differentiated Caco-2 cells (8 days post-confluence) intoxicated with TcdA and TcdB for 3, 6 and 8 hours. When compared to control cells treated with DMEM FBS-free alone, the immunostaining of unpermeabilized cells highlighted an increase in accessible Fibronectin (acc Fn) in the apical side of intoxicated cells, as depicted by the green fluorescence (**Fig. 1A**). Additionally, permeabilized cells were stained for total Fibronectin (total Fn) and nuclei (blue) (**Fig. 1A**). Representative confocal microscopy images, including 3D projections and magnified views, showcased an equal distribution of total Fn in both conditions (**Fig. 1A**) Interestingly, there is an evident accumulation of acc Fn around intoxicated cells when compared to control (**Figure 1A**). Quantitative analysis of relative fluorescent intensity (RFLi) showed no significant change in accessible and total Fn during intoxication (**Fig. S1A, Fig. 1C,D**). However, when observing the basal and apical side of cells individually, quantitative analysis showed that after 8 h of TcdA/TcdB-intoxication, there was a significant increase in acc Fn at the apical side of IECs (**Fig. 1C**). Yet, total abundance of Fn remains unchanged throughout the 8 hours of intoxication (**Fig. S1A**). In the basal side of intoxicated IECs, no significant change in accessible or total Fn was observed during the 8-hour intoxication (**Fig. 1D**). Altogether, these findings suggest that during intoxication, Fn is redistributed towards the apical side of IECs. Control western blots demonstrate that TcdA/TcdB-intoxication results in glucosylated Rac-1 (**Fig. 1G**).

**Fig. 1.**
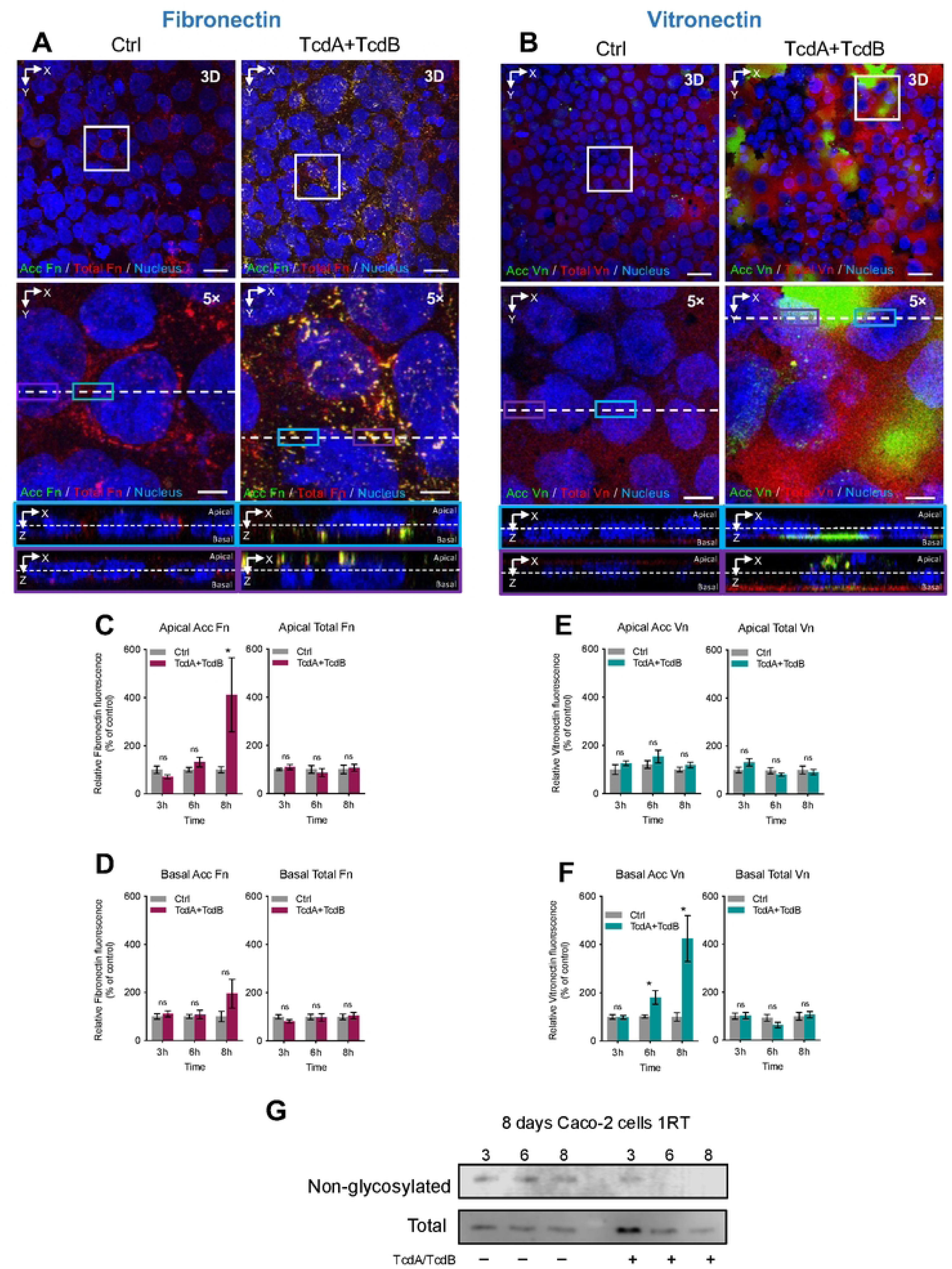
Effect of TcdA/TcdB intoxication of intestinal epithelial Caco-2 cells in redistribution of Fibronectin and Vitronectin. Differentiated Caco-2 cells intoxicated with TcdA and TcdB for 3, 6, or 8h in DMEM FBS-free. As a control, cells were treated with DMEM FBS-free. Unpermeabilized cells were stained for accessible Fibronectin or Vitronectin (acc Fn or acc Vn; green), permeabilized, and stained total Fibronectin or Vitronectin (total Fn or total Vn; red) and nuclei (blue). **a, b** Representative confocal microscopy images 3D projection of control cells (left) and intoxicated cells for 8h (right) immunostained for acc Fn or Vn and total Fn or Vn, below a magnified slide (XY), and the orthogonal view (XZ). Relative fluorescence intensity measured as the sum of raw intensity density/area for each z-step of accFn and total Fn, its abundance in the **c** apical side or **d** in the basal side of the cell; in the same way, the relative fluorescence intensity of Vn, its abundance in the **e** apical side and **f** the basal side of the cell. **g**, immunoblotting of anti- nonglucosylated Rac1 and total Rac1 of cell lysates of differentiated Caco-2 cells intoxicated with TcdA and TcdB for 3, 6, or 8 h. Nonglucosylated Rac1 was evaluated with corresponding antibodies, then the membrane was stripped, and subsequently tested for total Rac1. Western blotting is representative of 3 independent experiments. Controls were set at 100%. Error bars indicate the mean ± SEM from at least 9 fields (*n* = 3). Statistical analysis was performed by Two- Way ANOVA post-Bonferroni; ns, *p* > 0.05; * *p* < 0.05. Scale bar, top panels 20µm; bottom panels 5µm.

For Vn, we observed that TcdA/TcdB-intoxication of differentiated Caco-2 cells displayed distinctive changes in Vn distribution. In comparison to control cells, immunostaining of unpermeabilized cells revealed an increase in accessible Vn (acc Vn) on the basal side of intoxicated cells, as indicated by accumulation of the green fluorescence (**Fig. 1B**). Additionally, permeabilized cells were stained for total Vitronectin (total Vn) and nuclei (blue) (**Fig. 1B**).

Representative confocal microscopy images, along with 3D projections and magnified views, displayed equal distribution of total Vn surrounding the nucleus of the cells (**Fig. 1B**). Interestingly, there is an increase in green fluorescence which is accumulated in portions where cells have detached from the wells resulting in evident clusters of acc Vn (**Fig. 1B**). Quantitative analysis of RFLi showed a significant increase in accessibility but not in total Vn after 8 h of incubation (**Fig. S1B**). Further analysis shows that accessible Vn increase was on the basal side rather than apical, with increased of accessible basal levels of Vn in as early as 6 h post-intoxication (**Fig. 1E,F**). Despite these increments in Vn, no changes in total Vn were detected either in the apical or basal layer of monolayers of Caco-2 cells (**Fig. 1E,F**). These results suggest that changes in accessible Vn, are not due to increased apical levels of Vn, but due to detachment of Caco-2 intoxicated cells that lead to increased basal accessible vitronectin.

### TcdA/TcdB-intoxication of intestinal epithelial cells leads to increased accessibility of α_5_β_1_ and α_v_β_1_ integrins

In prior work, we demonstrated that *C. difficile* spores bind to Fn and Vn to gain intracellular access through their cognate integrin receptors, α_5_β_1_ and α_v_β_1_, respectively ^36^. Consequently, we also explored whether TcdA/TcdB-intoxication of IECs would lead to increased accessible α_5_β_1_ and α_v_β_1_ integrin receptor. Representative confocal micrographs show that there is an evident increase in accumulation of accessible α_5,_ but this was observed to happen in clusters around cells that seem to be surrounding potential sites of cellular apoptosis because of intoxication (**Fig. 2A**). Quantitative analysis unveiled a significant increase in RFLi of accessible ɑ_5_ (acc ɑ_5_) and total ɑ_5_ in cells intoxicated with TcdA/B compared to controls (**Fig. S2A,B**). Closer quantification of ɑ_5_ abundance in the apical side of IECs revealed a significant increase in accessible but not total ɑ_5_ integrin subunit (**Fig. 2D**). Conversely, a significant increase in accessible and total ɑ_5_ integrin was observed at the basal side of IECs after 8 h of toxin-intoxication (**Fig. 2E**).

**Fig. 2.**
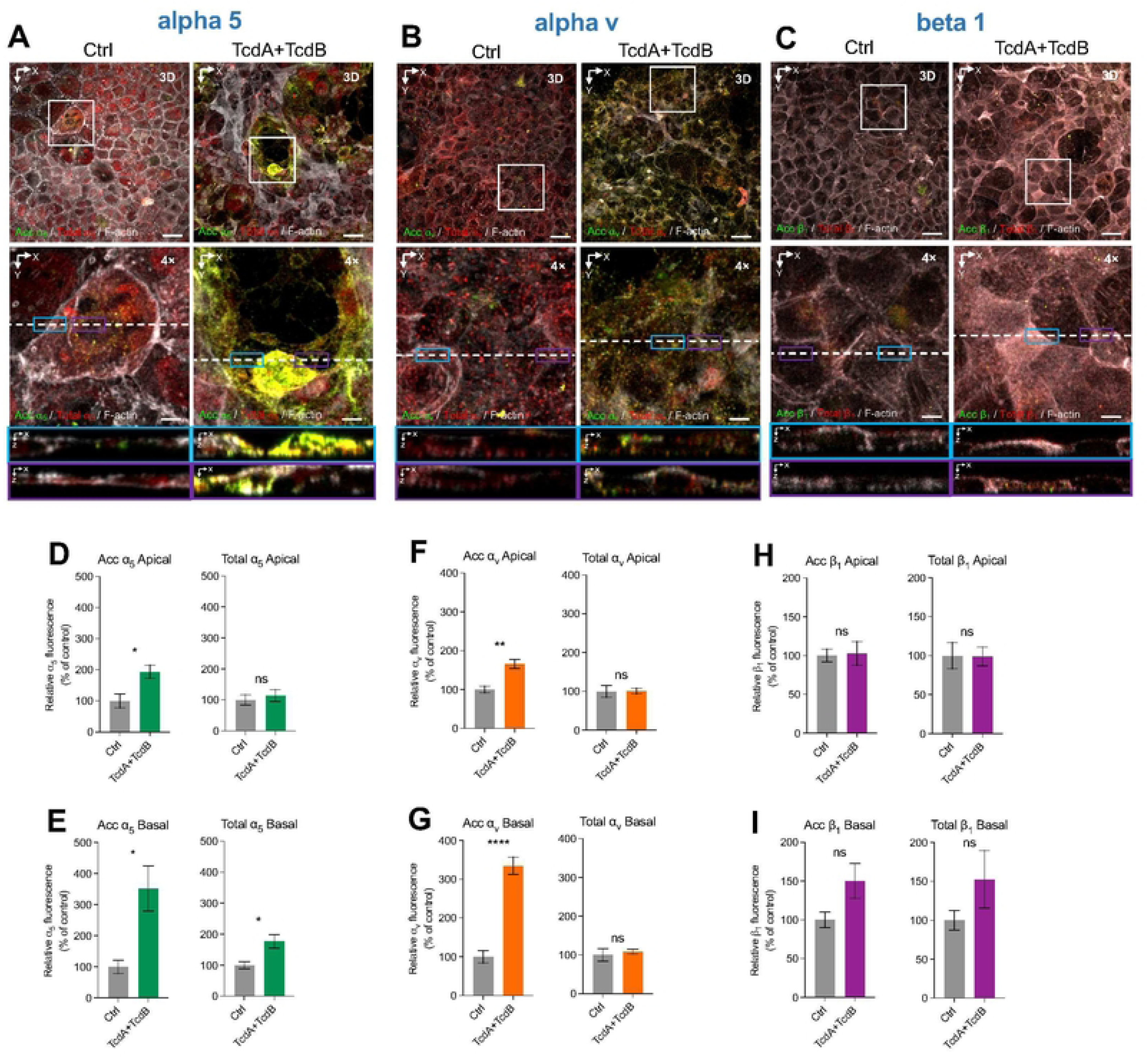
TcdA and TcdB increase accessible ⍺_5_ and ⍺_V_ but no β_1_ integrins in intestinal epithelial cells. Differentiated Caco-2 cells intoxicated with 600pM of TcdA and TcdB for 8h in DMEM FBS-free. As a control, cells were treated with DMEM FBS-free. Unpermeabilized cells were stained for accessible **a**, ɑ5 integrin; **b,** ⍺_V_ integrin, and **c**, β1 integrin, (shown in green), permeabilized, and stained total ɑ5, ⍺_V_ or β1 integrin respectively (shown in red) and F-actin (grey). **a-c**, Representative confocal microscopy images 3D projection of control cells (left) and intoxicated cells for 8h (right); below a magnified slide (XY), and the orthogonal view (XZ). **d-i,** Quantification of relative fluorescence intensity based on raw intensity density per area for each individual cell generated from the microscopy images using the 3D Surface Plotter plug-in of ImageJ. Relative fluorescence intensity measured as the sum of raw intensity density/area for each z-step of accessible and total ⍺_5_ located in the **d,** apical, and **e**, basal side of the cell. Relative fluorescence intensity measured as the sum of raw intensity density/area for each z-step of accessible and total ⍺_V_ located in the **f,** apical and **g**, basal side of the cell. Relative fluorescence intensity measured as the sum of raw intensity density/area for each z-step of accessible and total β_1_ located in the **h,** apical, and **i**, basal side of the cell. Controls were set 100%. Error bars indicate the mean ± S.E.M from at least 9 fields (*n* = 3). Statistical analysis was performed by unpaired Student’s *t* test, ns, *p* > 0.05; * *p* < 0.05; ** *p* < 0.01. Bars, top panels 20 µm; bottom panels 5µm.

For the integrin subunit α_v_, representative confocal micrographs show that there is an evident increase in accumulation of accessible α_v_ distributed equally in the observed tissue (**Fig. 2B**). Quantitative analysis revealed a notable increase in RFLi of accessible ɑ_V_ (acc ɑ_V_) but not total ɑ_V_ in cells intoxicated with TcdA/TcdB when compared to controls (**Fig. S2B**). Assessment of ɑ_V_ abundance in the apical side of IECs showed a significant increase in accessible but not in total ɑ_V_ integrin subunit (**Fig. 2F**). Similarly, a significant increase in accessible but not in total ɑ_V_ was observed on the basal side of IECs during intoxication (**Fig. 2G**).

In striking contrast to the redistribution of alpha-integrin subunits, no changes in the levels of accessible and total integrin subunit β_1_ was observed upon intoxication of Caco-2 cells with TcdA/TcdB (**Fig. 2C**). Representative confocal micrographs show that there is an equal distribution of β_1_ in both intoxicated and non-intoxicated cells (**Fig. 2C**). Quantitative analysis revealed no changes in the RFLi of accessible β_1_ (acc β_1_) as well as no significant change in total β_1_ levels in cells intoxicated with TcdA/B compared to controls (**Fig. S2C**). Examination of β_1_ abundance in the apical side of IECs revealed no significant changes in accessible and total β_1_ integrin (**Fig. 2H**). Likewise, there was no significant change in the abundance of accessible and total β_1_ integrin on the basal side of IECs during intoxication (**Fig. 2I**). These findings emphasize that TcdA/B doesn’t affect accessible or total β_1_ levels within IECs on both the apical and basal sides. Overall, these results indicate that TcdA/TcdB-intoxication of IECs increases accessible levels of ɑ_5_ and ɑ_V_ in both, apical and basal sides, while no change in accessible levels of β_1_ where observed. Intriguingly, TcdA/TcdB-intoxication increased the levels of total ɑ_5_ in the basal side, which is consistent with repair-functions of Fn-ɑ_5_ signaling^37^. Collectively, these data indicates that TcdA/TcdB-intoxication of IECs increases accessible ɑ_5_ and ɑ_V_ integrin receptors but does not affect availability of β_1_.

### Effect of TcdB-intoxication on the redistribution of intestinal Fn and Vn *in vivo*

To explore whether *C. difficile* toxin-intoxication would also lead to increase accessible Fn and Vn *in vivo*, we utilized an ileal ligated loop mouse model injecting various doses of TcdB (with 0.1, 0.5, 1 and 5 µg of TcdB) for 5 h prior to confocal analysis. Upon staining for Fn, we observed in non-intoxicated ileal loops sites with accessible and total Fn at various sites of the villus (**Fig. 3A and S3A**). Interestingly, holes at the dome of the villus, which appear empty in control samples, had a notorious increase in accessible Fn upon intoxication with 5 µg of TcdB (**Fig. 3A and Fig. S3A**). These holes are notoriously known as goblet cells as they do not stain for actin-cytoskeleton ^38^. However, upon analyzing how accessible and total fluorescence intensity per defined area changed upon TcdB-intoxication, we observed a slight increase in accessible and total fibronectin with 0.1 µg of TcdB (**Fig. 3C,D**), yet levels subsequently decreased by 40% when intoxicating with 5 µg of TcdB (**Fig. 3C,D**).

**Fig. 3.**
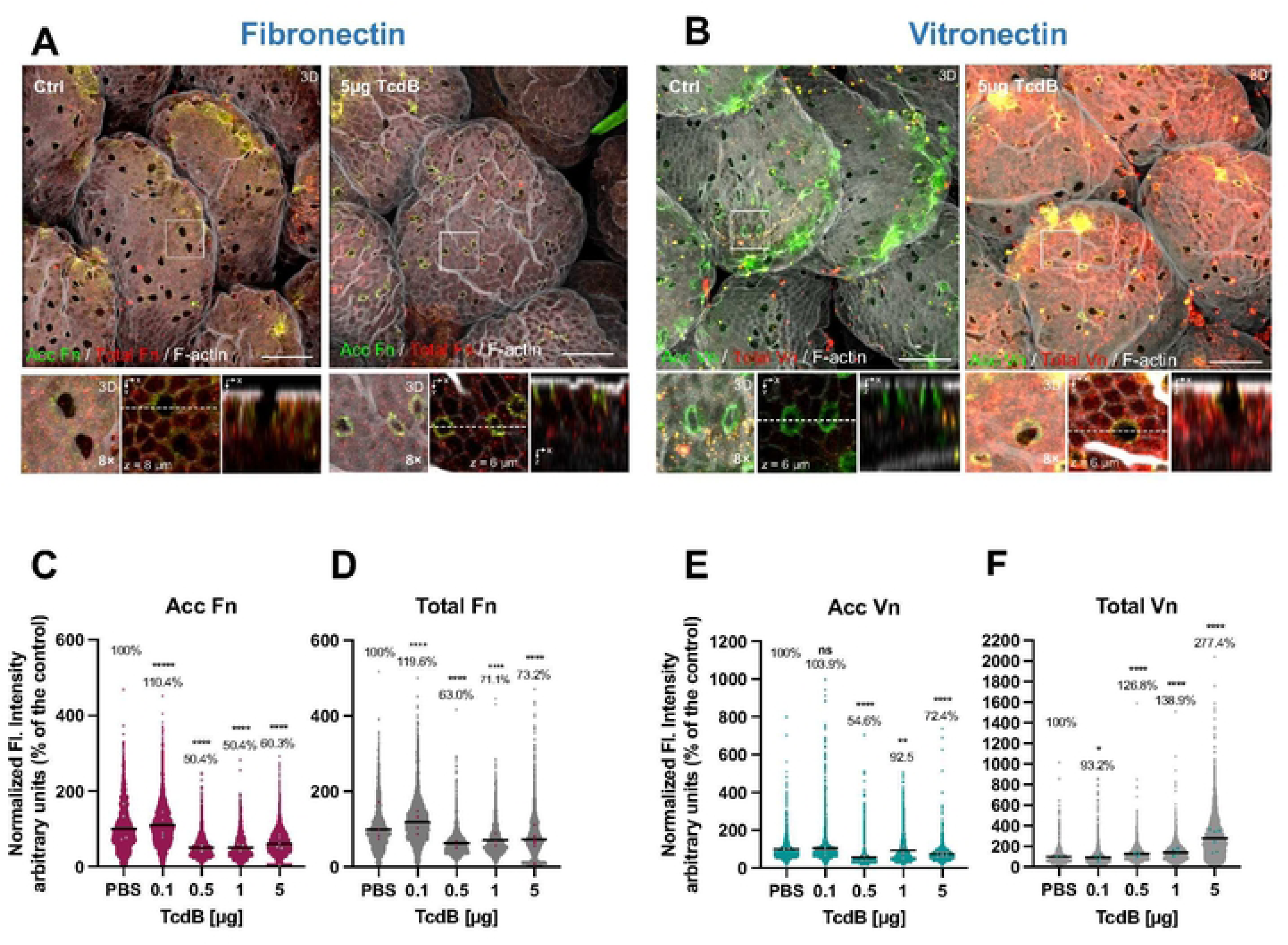
Effect of TcdB in the redistribution of Fibronectin and Vitronectin a ligated ileal loop mouse model. Ileal ligated loops were intoxicated for 5 h with 0.1, 0.5, 1, or 5µg of TcdB or saline as control. Then loops were removed, washed, fixed, and subjected to immunofluorescence. Unpermeabilized tissues were stained for accessible Fibronectin or Vitronectin (acc Fn or acc Vn; green), and then permeabilized and stained total Fibronectin or Vitronectin (total Fn or total Vn; red) and F-actin (grey). **a**-**b**, Representative confocal microscopy images 3D projection of control cells (left) or intoxicated loops with 5µg TcdB (right) immunostained for accessible and total Fn or Vn; right bottom, a magnified 3D projection, next to a z-stack (XY), and then magnified orthogonal view (XZ). Quantification of **c, e**, accessible or **d, f**, total Fn or Vn fluorescence intensity per cell measured in the z-projection (sum). For acc Fn or Vn, the analyzed area was Ctrl of 170,360 µm^2^; 0.1 µg TcdB of 340,720 µm^2^; 0.5 µg TcdB of 340,720 µm^2^; 1 µg TcdB of 340,720 µm^2^ and 5 µg TcdB of 511,080 µm^2^. n = 3 animal per group. In scatter plots, each dot corresponds to one independent cell. Dots in colors correspond to the average of each analyzed mice/field. Error bars indicate mean or mean ± SEM. Statistical analysis was performed by unpaired Student’s *t* test; ns, *p* > 0.05; * *p* < 0.05; \*\**p* < 0.01; **** *p* < <0.0001. Scale bar 20 µm.

In contrast to Fn redistribution, healthy un-intoxicated ileal tissue had significant accessible Vn in various sites in the villi surface, and particularly in goblet-like cells (**Fig. 3B and Fig. S3B**). Strikingly, upon intoxication, no qualitative changes where evident with 0.5 or 1.0 µg of TcdB (**Fig. S3B**). However, intoxication with 5 µg of TcdB led to a massive increase in Vn throughout the entire intestinal epithelial layer, and this increase was primarily in total intracellular Vn (**Fig. 3B**). Upon quantitative analysis of changes in Vn, while accessible Vn decreased in a TcdB- concentration dependent manner (**Fig. 3E**), we observed a striking TcdB-concentration dependent increase in total Vn was observed, with a ∼ 3-fold increase in the presence of 5 µg of TcdB (**Fig. 3F).** In summary, these results indicate that TcdB-intoxication of intestinal tissue *in vivo* leads to overall decrease in accessible and total Fn, accessible increase is evident in goblet-like cells. By contrast, TcdB-treatment while decreasing accessible Vn, led to substantial increase in total intracellular Vn.

### Impact of *C. difficile* toxin-intoxication in spore-Fn and -Vn associations

Despite *in vitro* work showing that Fn and Vn interact with *C. difficile* spores with strong *K*_D_,^39^ and that these molecules increase during TcdA/TcdB-intoxication of IECs as shown in **Fig. 1A,B**, it is unclear whether intestinal Fn and Vn interacts with *C. difficile* spores. First, we tested whether TcdA/TcdB-intoxicated IECs release Fn and Vn to bind to *C. difficile* spores. For this, *C. difficile* spores were incubated with DMEM (control), supernatant of untreated or 8 h TcdA/TcdB-treated differentiated monolayers of Caco-2 cells. Upon immunostaining of *C. difficile* spores for Fn and Vn, while no immunoreactivity was observed in DMEM-incubated spores, significant Fn ad Vn were detected in spores incubated with untreated supernatant (**Fig. 4A,B**). Notably, TcdA/TcdB-intoxication lead to a significant increase in supernatant Fn and Vn binding to *C. difficile* spores (**Fig. 4A,B**). Single spore quantification of Fn and Vn binding confirms these results and demonstrate a nearly two-fold increase in Fn and Vn binding when comparing supernatants from untreated cells with TcdA/TcdB-intoxicated cells (**Fig. 4C,D**). These results suggest that *C. difficile* spores might interact with intestinal Fn and Vn in an intoxicated mucosal layer.

**Fig. 4.**
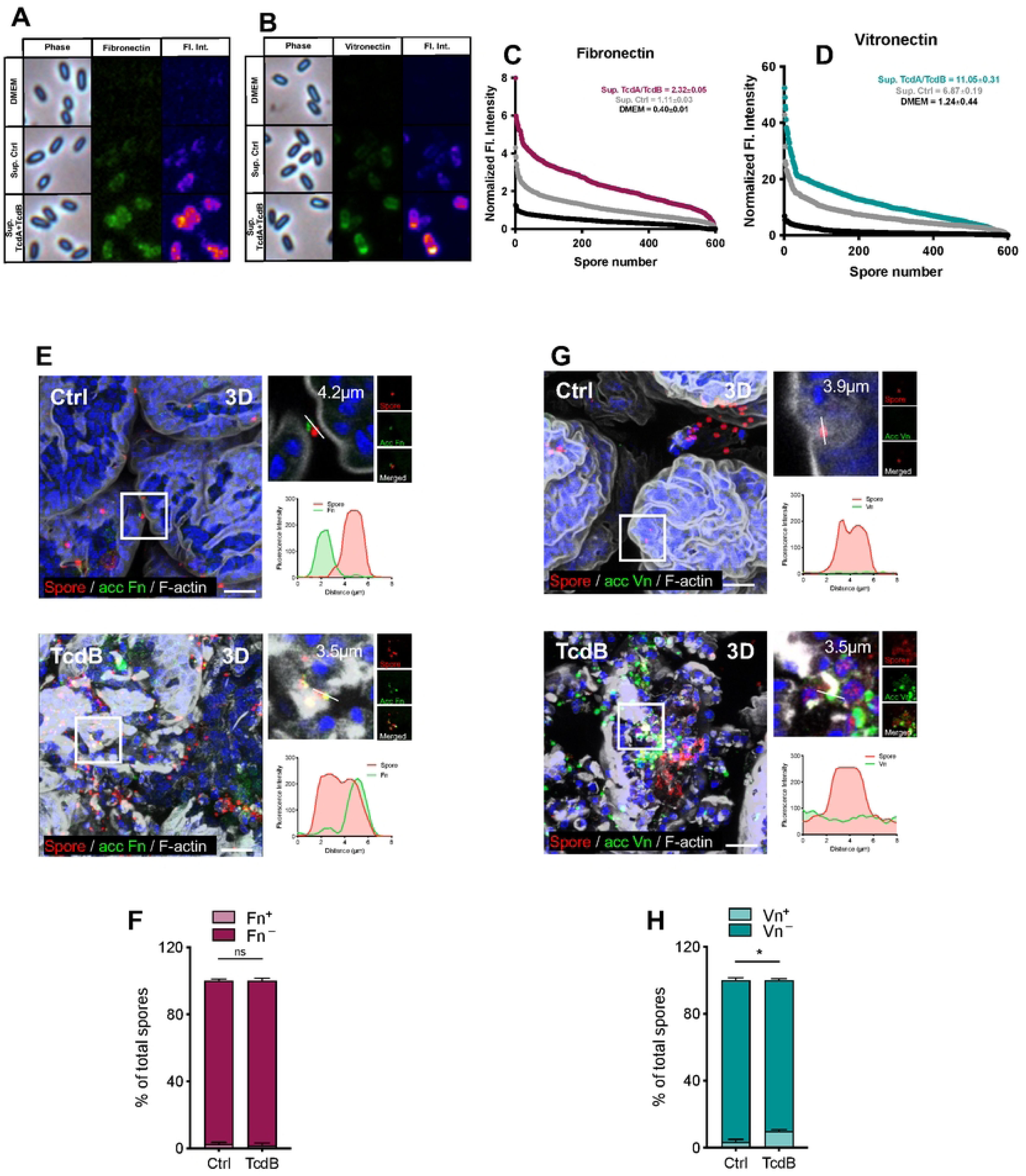
Intoxication increases association of *C. difficile* spores with Fn and Vn. a-d, Differentiated Caco-2 cells were intoxicated with 600pM of TcdA and TcdB for 8h in DMEM FBS-free media. Controls include non-intoxoicated cells in DMEM FBS-free. Supernatant was collected from untreated and intoxicated cells and subsequently utilized to resuspend *C. difficile* spores and incubate for 1 h at 37 °C. Spores resuspended in DMEM alone were also included as a control. Spores were washed and strained for immunofluorescence anti- fibronectin and - vitronectin. **a-b,** Micrographs show representative phase-contrast (phase), fibronectin and vitronectin specific immunofluorescence and fluorescence intensity profiles (Fl. int.). Representative Fl. int. were provided using 3D Surface plotter function of Fiji. **c**-**d**, Quantitative analysis of the fluorescence Fl. int. of Fn and Vn in spores Fl. Int of 600 spores. Mean ± SEM are denoted. **e**-g, Ileal ligated loops were intoxicated with 5µg of TcdB and 5 × 10^8^ *C. difficile* R20291 spores for 5 h. Then loops were removed, washed, fixed, and subjected to immunofluorescence. Unpermeabilized tissues were stained for accessible Fibronectin or Vitronectin (acc Fn or acc Vn; green), and then permeabilized and stained total Fibronectin or Vitronectin (total Fn or total Vn; red) and F-actin (grey). **e, g**, representative 3D confocal micrograph projection reconstruction of fixed whole-mount small intestine tissue, and magnification of *C. difficile* spores associated with Fn or Vn. Plot profiles of fluorescence intensity of *C. difficile* spores (red line) and accessible Fn or Vn (green lines). **f,h**, quantification of spores that were positive (Fn+) or negative (Fn-) for Fn fluorescence signal in **f**, or positive (Vn+) or negative (Vn-) for Vn fluorescence signal in **h**. The average of associated and non-associated spores with **f** Fn or **h** Vn for each field. A total of ∼ 500 spores were counted per mice (n = 5 per group). GRUBB’s test was performed to identify outliers, and one point was removed in Vn. Error bars indicate mean ± S.E.M. Statistical analysis was performed by two-tailed unpaired Student’s t test; ns indicates non-significant differences. Scale bar, 20 μm.

To explore the hypothesis that *C. difficile* spores interact *in vivo* with intestinal Fn and Vn, ileal loops injected with 5 µg of TcdB and 5 x 10^7^ spores, incubated for 5 h prior to immunostaining for accessible Fn and Vn and for total *C. difficile* spores. In the absence of TcdB, very few of adhered *C. difficile* spores (∼ 2 % of total spores) interacted with Fn (**Fig. 4E,F**). In striking contrast to *in vitro* results (**Fig. 4A,C**), TcdB-intoxication had no impact in the fraction of *C. difficile* spores associated with Fn, which remained similar to that of untreated ileal loops (**Fig. 4E,F**). The analysis of *C. difficile* spore association with Vn revealed significant differences between unintoxicated and TcdB-intoxicated ileal loops. In unintoxicated ileal loops, approximately 3% of total spores were found to be associated with Vn(**Fig. 4G,H**). However, this percentage increased markedly in TcdB-intoxicated ileal loops, where about 10% of total spores showed association with Vn. This increase was statistically significant, indicating a notable effect of TcdB intoxication on the interaction between *C. difficile* spores and Vn (**Fig. 4G,H).** Collectively, these results support the notion that *C. difficile* toxin-intoxication leads to increase association of *C. difficile* spores mainly with intestinal Vn, and to a lesser extent with intestinal Fn.

### TcdB increases *C. difficile* spore adherence and internalization to intestinal epithelial cells

In prior work, we demonstrated that toxin-intoxication of Caco-2 cells leads to increase spore adherence and internalization^33^, yet whether this also occurred *in vivo* remains unclear. Therefore, to address this question, ileal ligated loops were injected with 5 × 10^8^ *C. difficile* R20291 spores and 1 μg, or 5 μg of TcdB for 5h, or saline alone as control. Representative confocal images indicate a notable spore accumulation following intoxication, with clusters displaying a distinct affinity for specific regions within the cells, rather than an even distribution (**Fig. 5A**). Quantitative analysis of the number of spots (spores) adhered per 10^5^ μm^2^ relative to the control or internalized spores in the ileum in relation to the total spores (**Fig. 5B,C**), showed that while no changes in adhered and internalized spores where observed during intoxication with 1 μg of TcdB, a there was a nearly 4-fold increase in adhered spores (**Fig. 5B**), and 10-fold increase in internalized spores (**Fig. 5C**) were observed in the presence of 5 μg of TcdB. Overall, these results confirm that *C. difficile* toxin-intoxication of the intestinal epithelial layer increases adherence and internalization of *C. difficile* spores *in vivo*.

**Fig. 5.**
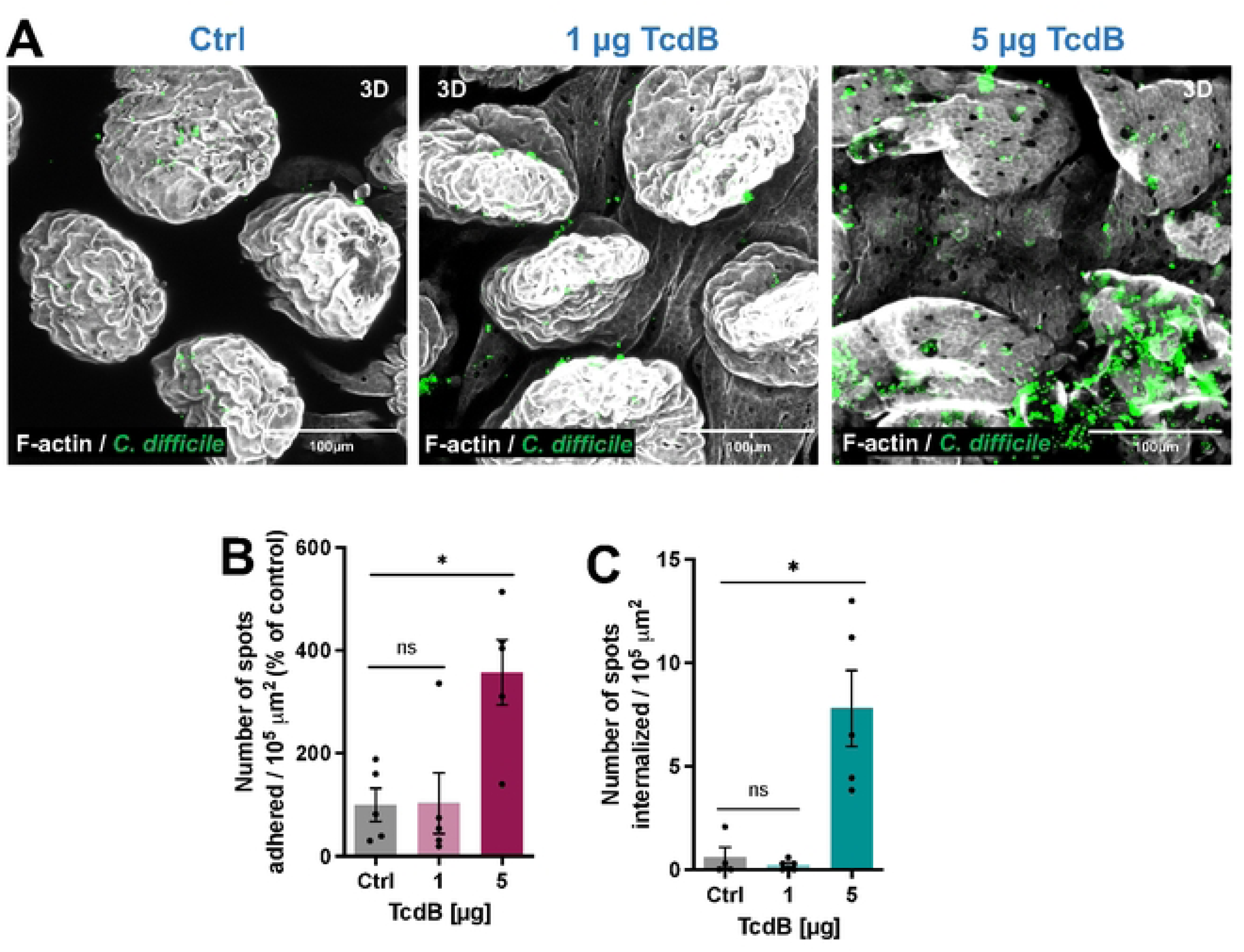
TcdB-intoxication increases *C. difficile* spore adherence and internalization to the intestinal barrier *in vivo*. Intestinal loops, of approximately ∼1.5 cm, were injected with 5 × 10^8^ *C. difficile* R20291 spores and 1 or 5 μg of TcdB for 5h (or saline alone as control). **a**, 3D projections of representative confocal micrographs. *C. difficile* spores are shown in green, and F- actin is shown in grey (fluorophores colors were digitally reassigned for a better representation). **b,c,** Quantification of **b** adhered or **c** internalized spots (spores) per 10^5^ μm^2^ is expressed in relative values to the unintoxicated control. Controls were set 100%. GRUBB’s test was performed to identify outliers, and one point was removed in **b** for group intoxicated with 1 μg TcdB and in **c** for control group. Error bars indicate the mean ± S.E.M. Statistical analysis was performed by unpaired Mann-Whitney test, ns, indicates non- significant differences, * *p* < 0.05. Scale bar 100μm. n = 5 mice per group.

### Protective effect of Bezlotoxumab in the redistribution of fibronectin and vitronectin during intoxication of intestinal epithelial cells

Clinical data demonstrates that administration of Bezlotoxumab correlates with a reduction in the rates of recurrent CDI ^40, 41^. In prior work, we demonstrated that Fn and Vn contributes to spore-internalization into IECs, and that intracellular spores are implicated in CDI recurrence ^36^. Hence, we sought to assess the impact of Bezlotoxumab in the TcdB-intoxication mediated redistribution of Fn and Vn in the intestinal mucosa. Although the reported dose utilized in mice has been reported to be ∼50 mg/kg^29, 42^, in our preliminary results we found that for our batch of Bezlotoxumab that dose proved to be lethal upon intraperitoneal injection, while 5 mg/kg yielded 100% survival (data not shown). Consequently, 5 mg/kg of Bezlotoxumab was utilized in downstream experiments. For this, 24 h prior to ileal loop surgery, mice were given an intraperitoneal injection of 5 mg/kg Bezlotoxumab or of saline (PBS). Next, ligated loops where injected either PBS (control) or 5µg TcdB. Additionally, representative confocal micrographs show that accessible Fn seems to decrease upon intoxication (**Fig. 6A**), similarly as in **Fig. 3A**, this trend is also observed in the presence of Bezlotoxumab (**Fig. 6A)**. Upon quantitative analysis of abundance of Fn, a significant decrease in accessible and total Fn was observed upon intoxication with TcdB. While presence of Bezlotoxumab in intoxicated ileal loops did not impact the levels of accessible when compared to intoxicated ileal loops alone, it caused a significant reduction in total Fn to ∼25 % of total when compared to unintoxicated ligated loops (**Fig. 6C**).

**Fig. 6.**
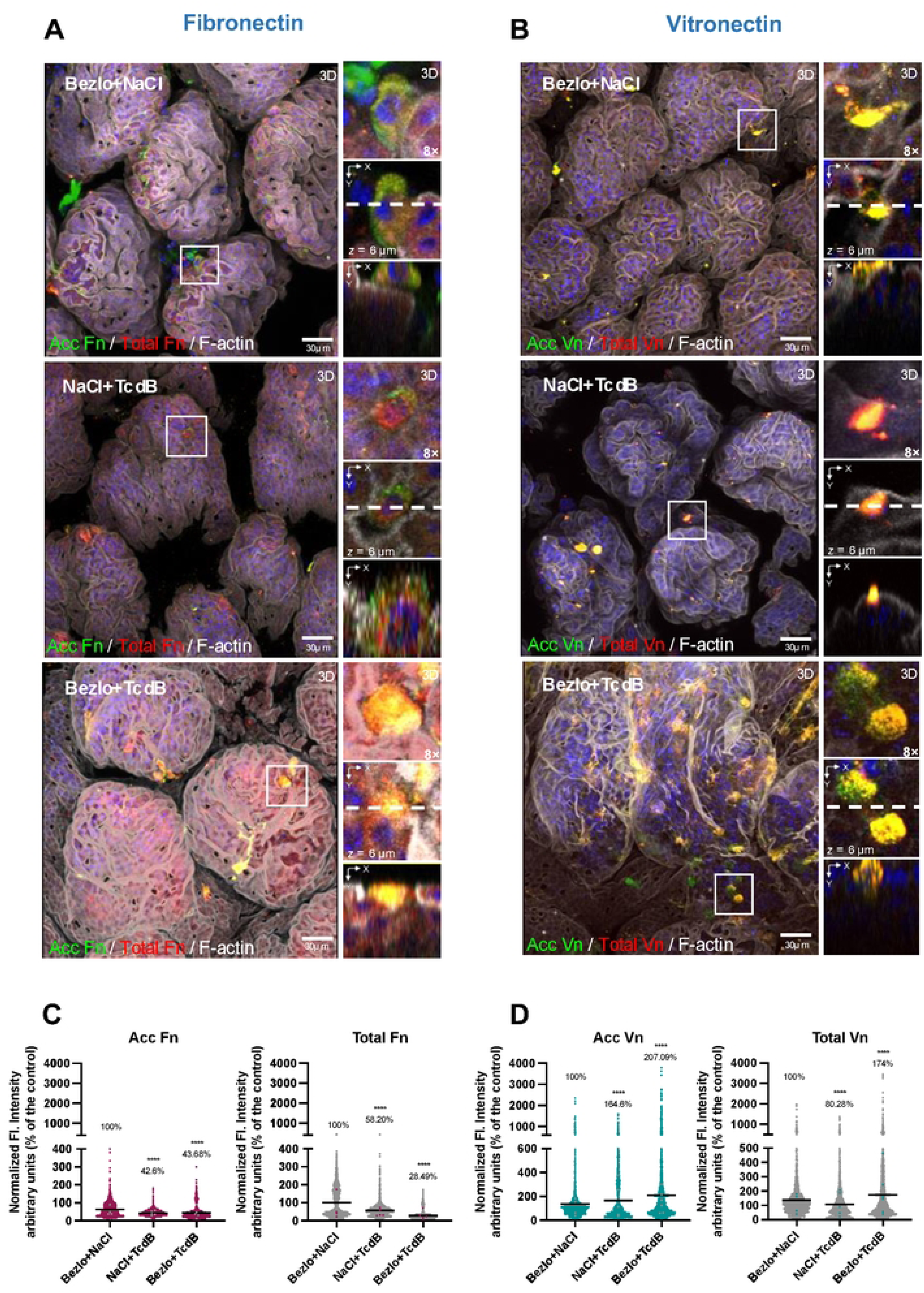
Effect of Bezlotoxumab in TcdB-intoxication induced redistribution of Fibronectin and Vitronectin *in vivo*. Prior to ileal surgeries intraperitoneal injection of 5mg/kg of Bezlotoxumab or saline solution was administered as indicated. Ligated loops were injected with *C. difficile* Spores and 5µg of TcdB or saline as control. Mice were let for recovery for 5 h before euthanasia. **a,b** Representative 3D confocal micrograph projection reconstruction of fixed whole-mount small intestine tissue. Unpermeabilized tissues were stained for Acc Fn or Vn shown in green, and then permeabilized and stained for total Fn or Vn shown in red, and F-actin in gray and cell DNA with Hoechst (blue), (some fluorophores colors were digitally reassigned for a better representation). Panels **c,d** show the fluorescence intensity immunodetected in cells for acc and total **c** Fn or **d** Vn. Nearly 2000 – 3000 cells were counted per field. Quantification of total or accessible Vn, fluorescence intensity per cell measured in the z-projection (sum). Error bars indicate mean ± S.E.M. Statistical analysis was performed by two-tailed unpaired Student’s t test, ns indicates non-significant differences; * *p* < 0.05; \*\**p* < 0.01; **** *p* < <0.0001. Colored dots represent the average normalized fluorescence intensity of each independent mice in the group. N = 5 mice per group. Scale bar, 30 μm.

In the case of Vn, representative confocal micrographs show that cell-types with increased accessible and total Vn became evident in TcdB-intoxicated ligated loops when compared to Bezlotoxumab treatment alone (**Fig. 6B**). Notably, intoxication of ligated loops in the presence of Bezlotoxumab lead to a substantial increase in these cell types expressing increased accessible and total Vn (**Fig. 6B**). Quantification analysis of the fluorescence intensity of accessible and total Vn reveals a significant increase in accessible Vn, but not total, in TcdB-treated ligated loops (**Fig. 6D**). Strikingly, in TcdB-intoxicated ligated loops in the presence of Bezlotoxumab, both, accessible and total Vn increased significantly (**Fig. 6D**), consistent with some cells having higher levels of accessible and total Vn. Taken together, these results indicate that while administration of Bezlotoxumab has no impact in levels of accessible and total Fn during TcdB-intoxication, it dramatically increases both accessible and total Vn in a subpopulation of intestinal epithelial cell types.

### Bezlotoxumab reduces spore adherence in TcdB-intoxicated mucosal epithelium layer

The increase in adhered and internalized *C. difficile* spores in a TcdB-intoxicated ligated loop, led us to hypothesize that prior administration of Bezlotoxumab could protect from increased adherence and internalization of spores into IECs *in vivo*. For this, we treated mice with an *i.p.* of 5 mg/kg of Bezlotoxumab or saline (NaCl) 24 h before surgery. Ligated loops where then injected with *C. difficile* spores in the presence or absence of 5 µg of TcdB (n = 4-5 per group). Interestingly, representative confocal micrographs show that during TcdB-intoxication, *C. difficile* spores adhered to IECs in clusters, as opposed to what was observed in the control treated with Bezlotoxumab alone (**Fig. 7A**). Upon analysis of micrographs of TcdB-intoxicated and Bezlotoxumab-treated ligated loops, there was an evident decrease in the levels of spores associated with the intestinal mucosa in the Bezlotoxumab treated tissue(**Fig. 7A**). Upon performing quantitative analysis, we observed that in tissues intoxicated with TcdB, spore adherence increases significantly by ∼400 % when compared to Bezlotoxumab control (**Fig. 7B**). In animals pre-treated with Bezlotoxumab and intoxicated with TcdB, spore adherence increased significantly by 280 % when compared to Bezlotoxumab control (**Fig. 7B**). Quantification of intracellular spores demonstrated that while there was an increase in intracellular spores in TcdB-intoxicated ileal loops, which decreased in the presence of Bezlotoxumab, these changes where not significant (**Fig. 7C**). Collectively, these results indicate that Bezlotoxumab might prevent TcdB-mediated enhanced adherence and internalization into the intestinal mucosa.

**Fig. 7.**
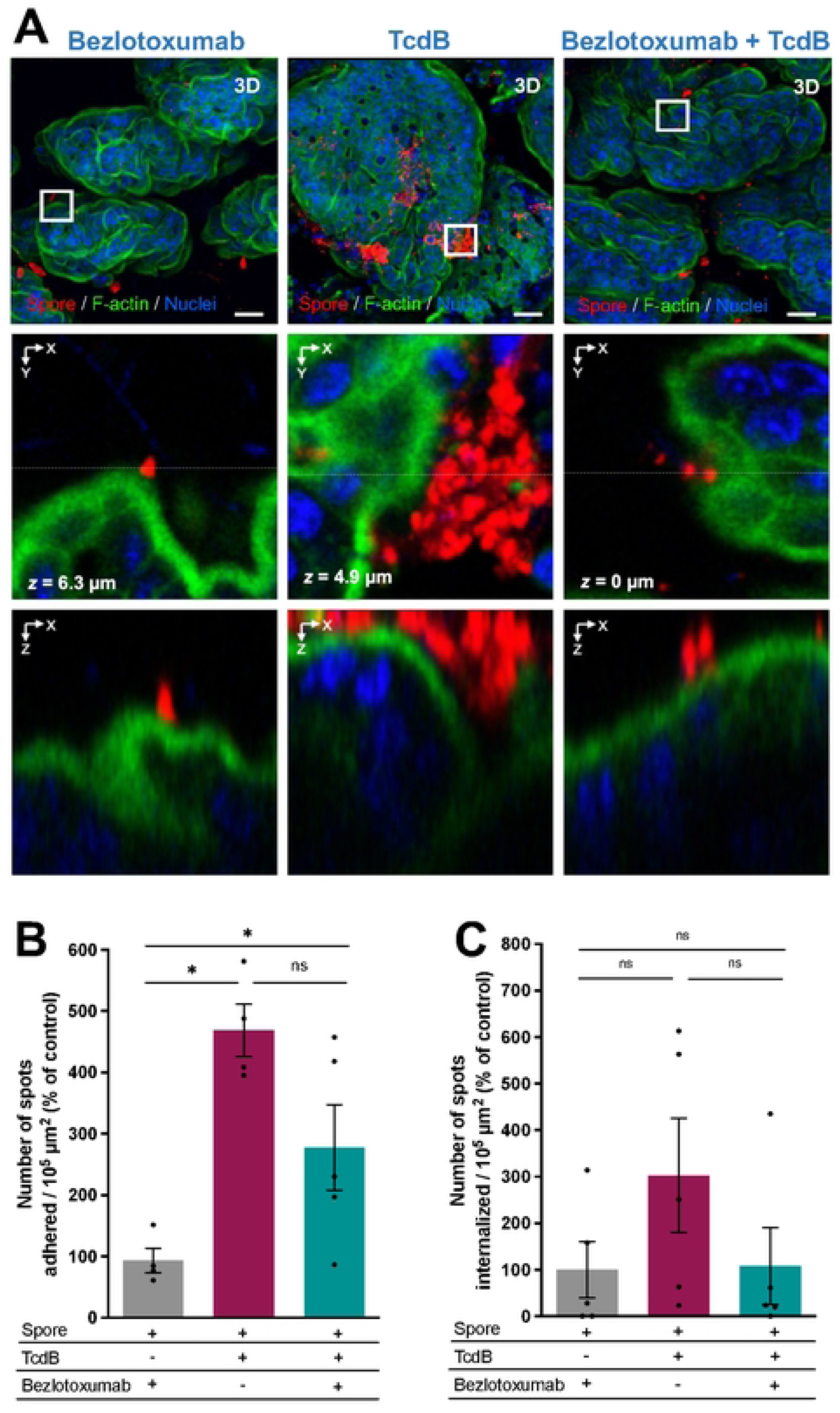
Bezlotoxumab neutralizes TcdB- increased adherence of *C. difficile* spores caused by TcdB. **A)** Mice were treated with an intraperitoneal injection of 5mg/kg of Bezlotoxumab or saline solution 24h prior to ileal loops surgery. Loops where then injected with 5 × 10^8^ *C. difficile* R20291 spores with or without 5μg of TcdB. Representative confocal micrographs of ileal loops are shown, with *C. difficile* spores stained in red, F-actin is shown in green, and nuclei in blue. **b,c,** Quantification of **b** adhered and **c** internalized spots (spores) per 10^5^ μm^2^ is shown relativized to the non-intoxicated Bezlotoxumab control. GRUBB’s test was performed to identify outliers, and one point was removed in group Bezlotoxumab + spore and one in TcdB + spore. Error bars indicate the mean ± S.E.M. Statistical analysis was performed by Mann-Whitney test, ns, *p* > 0.05, * *p* < 0.05. Scale bar 20μm. n = 5 mice per group.

## Discussions

During CDI, toxin production and spore formation play crucial roles in disease manifestation and recurrence, respectively. ^27, 43^. Recent research has significantly advanced our understanding of how *C. difficile* interacts with host cells ^33, 36^. *C. difficile* spores utilize E-cadherin for adherence and Fn and Vn integrin-dependent pathways for internalization into IECs^33, 36^, with bothspore-adherence and -internalization contributing to disease recurrence^36^. However, the impact of *C. difficile* toxins in spore adherence and internalization remains unclear. This study reveals that *C. difficile* toxin-intoxication of IECs contributes to remodeling of Fn, Vn, and their cognate integrins. Specifically, TcdA/TcdB intoxication led to increased binding of *C. difficile* spores with released Fn and Vn from Caco-2 cell monolayers, but only with Vn in intoxicated ligated loops. Importantly, TcdB-intoxication increased spore adherence and internalization into intestinal tissue. Using Bezlotoxumab, a monoclonal antibody that neutralizes TcdB, we observed that neutralization of TcdB had no impact on redistribution of Fn and Vn, however there was a reduction in spore adherence and internalization into IECs *in vivo*. These findings provide valuable insights into the role of *C. difficile* toxins TcdA and TcdB in spore interactions with the intestinal tissues.

Our work expands our understanding of the roles of *C. difficile* major toxins, TcdA and TcdB, beyond their essential function in causing clinical symptoms and inflammation during infection.^8^. Our data suggests that TcdA/TcdB-intoxication enhances the levels of accessible bioavailable Fn and Vn from human IECs. While Fn and Vn are primarily located in the basal and basolateral membrane of IECs and contribute to their polarity,^11, 12^ accessible Fn and Vn are commonly identified in certain epithelial folds and extrusion sites that undergo adherent junction reorganization ^36, 44–46^.

Both TcdA and TcdB glucosylate host Rho family of GTPases ^47^; which blocks the exchange of GDP for GTP and prevents RhoA, Rac1 and Cdc42 from functioning. This leads to cell rounding, opening of tight and adherent junctions and cell death ^48^. The toxin-mediated cell- intoxication likely results in a remodeling of the basolateral membrane, increasing accessibility of Fn and Vn and causing extrusion of Caco-2 cells. This is supported by the observed increase in apical and basal levels of the α_5_ and α_v_ integrins. Both toxins also elicit a robust proinflammatory response through multiple pathways, including mitogen-activated protein kinases that activate the transcription nuclear factor κB (NF-κB)^49, 50^. They also trigger the ASC-containing inflammasome, leading to secretion of various cytokines such as IL-8, tumor necrosis factor alpha (TNF- α), and IL-6, and IL-1B ^51–53^. Additionally, these toxins also trigger a series of cell death pathways including apoptosis, necrosis and pyknosis ^54–57^. Under the experimental conditions tested, the observed changes in levels of accessible and total Fn and Vn are likely due to a combination of intoxication, rupture of tight and adherent junctions and to some extent to cell death and detachment. Prior work by Schwan et al. demonstrated that TcdB, but not TcdA, caused a slight change in apical Fn ^25^. However, the *C. difficile* binary toxin rapidly reroutes Fn-containing vesicles from the basal to the apical side of Caco-2 cell monolayers within 1 hour of intoxication ^25^. This Fn redistribution occurs through binary toxin-induced protrusions, primarily involving Rab11 vesicle traffic to the apical surface, colocalizing Fn with α_5_ and β_1_ integrins ^25^. Notably, accumulations of laminin and Vn were also observed, albeit to a lesser extent than Fn ^25^.

Given the extended time (∼8 hours) required in our study to observe these changes, it is likely that alterations in accessible Fn and Vn are due to secondary cell death and/or opening of the basolateral junctions, rather than a specific pathway. However, early work shows that DSS-induced inflammation of IECs also leads to increased Fn production ^22^. Therefore, it is plausible that TcdA/TcdB toxin-mediated inflammation also leads to increased Fn production. Further work to elucidate the specific signaling pathways involved in the increased Fn and Vn levels, as well as the individual roles of each toxin is warranted.

This work also demonstrates that, at least TcdB-intoxication of intestinal tissue increases total gut Vn in intestinal tissue. Surprisingly, in contrast to the increase of both accessible Fn and Vn in Caco-2 cells, only total Vn levels changed during TcdB-ileal loop intoxication. This discrepancy might reflect the differences between experimental platforms and/or additional pathways activated in the different environments. The inherent higher variability of immunofluorescence of these molecules in ligated loops compared to tissue monolayers could be a limitation of our technique. Despite these inconsistencies, the finding that TcdB-intoxication leads to an increase in total Vn is novel. Confocal micrographs clearly show that the main source of total Vn was intracellular, primarily in the monolayers of IECs. Our understanding of gut Vn is limited, but it is known to be produced by IECs during early stages of development ^58^, and its increased expression during chronic intestinal inflammation has been linked to protection against colitis ^24^.

Both TcdA and TcdB toxins are known to induce inflammasome activation^51, 53^, but whether this impacts intracellular Vn production remains unclear and is a subject of ongoing research in our lab. Although TcdA and TcdB have similar functional domains and cause comparable changes in collapsing tight and adherent junctions and in cell physiology, recent studies show that they distinctively alter host pathways^59^. For instance, TcdB, but not TcdA, affects stem cells, impacting tissue repair and disease recurrence prevention^60^. TcdA and TcdB toxins also have different receptor requirements. TcdA binds to glycoproteins like sucrose isomaltose (SI) and glycoprotein 96 (gp96) ^61, 62^, sulfated glycosaminoglycans (sGAGs) and/or membrane low-density lipoprotein receptor (LDLR) family ^63^, with direct binding to LRP1 ^64^. In contrast, TcdB has three reported protein receptors: chondroitin sulfate proteoglycan 4 (CSPG4), Frizzled 1 (FDZ1), FZD2, FZD7 and Nectin 3 (also called “polivirus receptor-like 3” or PVRL3) ^65–67^. The distribution of these receptors in the murine GI tract remains unclear, but it’s possible that TcdA also impacts Vn accessibility, and perhaps Fn. Furthermore, toxin-mediated epithelial cell death initiates a MAP kinase signaling cascade, triggering recruitment and infiltration of granulocytes and activation of innate immune cells. This complex interplay between toxins, host cells, and immune responses contributes to the pathogenesis of *C. difficile* infection and highlights the need for further research into the specific mechanisms involved.

Previous work demonstrated that Fn and Vn bind to *C. difficile* spores in a concentration-dependent manner through solid-phase binding and pulldown assay^39^. This current study expands on those findings by showing that *C. difficile* spores bind to Fn and Vn released from TcdA/TcdB-intoxicated cells. Additionally, we observed increased spore-Vn association in TcdB-intoxicated intestinal loops, proving the first evidence of *C. difficile* spores associating with Vn *in vivo*. This aligns with the increased levels of total gut Vn observed in TcdB-intoxicated intestinal tissue.

The specific spore surface ligand responsible for Fn and Vn binding remains unclear, with no obvious candidates identified in the *C. difficile* spore exosporium proteome ^68^, current matter of investigation in our lab. In contrast, several Fn-binding proteins have been identified on the surface of *C. difficile* vegetative cells. *C. difficile*’s genome encodes at least two cell-surface proteins for which experimental evidence supports their role as Fn-binding proteins (i.e., FbpA and Fbp68) ^69, 70^, and at least FbpA is required for normal colonization of germ-free mice ^69^. While we have previously demonstrated that *C. difficile* spores use Fn and Vn as molecular bridges to internalize into human intestinal epithelial Caco-2 cells^36^, the implications of these interactions during *C. difficile* infection, particularly in terms of persistence and disease recurrence, remain to be fully elucidated.

A major conclusion provided by this work is that TcdB-intoxication of intestinal tissue enhances the adherence and internalization *C. difficile* spores *in vivo*. This finding extends previous research that demonstrated that TcdA/TcdB-intoxication of Caco-2 cells monolayers led to higher levels of spore adherence and internalization^33^, by showing that this phenomenon also occurs *in vivo*. Although the study focused on TcdB, substantial evidence suggests that both toxins A and B, significantly disrupt the epithelial barrier ^71–73^, indicating that these results may also extrapolate to TcdA. Previous work showed that TcdA/TcdB-intoxicated monolayers of Caco-2 cell monolayers underwent adherent junction remodeling, increasing accessible E-cadherin and its association with *C. difficile* spores ^33^. Moreover, E-cadherin was identified as a spore-adherence receptor that contributed to spore entry into IECs.

While changes in accessible E-cadherin were not quantified in this work, this and additional mechanisms, including enhanced interactions between *C. difficile* spores and intestinal Vn, may contribute to increased adherence and internalization. These results also support the hypothesis that during infection, *C. difficile* toxins A and B contribute to intestinal epithelial remodeling, potentially increasing spore persistence and, consequently, disease recurrence after vancomycin treatment. Our lab is currently investigating the specific impact of each toxin in *C. difficile* persistence and disease recurrence.

A final contribution of this work pertains the impact of Bezlotoxumab on TcdB-mediated remodeling of Fn and Vn, and consequently, on adherence and internalization of *C. difficile* spores to intestinal tissue. Initial clinical trials by Merck, which included anti-TcdA and anti-TcdB monoclonal antibodies (i.e., Actoxumab and Bezlotoxumab), showed that Bezlotoxumab significantly reduced CDI recurrence when combined with antibiotics ^41, 74, 75^. In preclinical animal models, both, Actotoxumab and Bezlotoxumab together protect mice from severe *C. difficile* disease ^76, 77^. However, these previous studies used an *i.p.* dose of 50 mg/kg ^76, 77^, which is 10-fold higher than the dose used in this study (5 mg/kg). In preliminary work, we attempted to use 50 mg/kg, however, this dose proved lethal in our murine model system. Consequently, we identified 5 mg/kg as a non-lethal dose that could be administered via *i.p.* 24 h prior to the ileal loop surgery. Our data suggests a trend in preventing TcdB-mediated adherence and internalization into intestinal tissue, but higher levels of Bezlotoxumab might be necessary to fully prevent this effect. Similarly, TcdB-mediated increased total gut Vn did not decrease upon administration of Bezlotoxumab. Overall, while our data strongly supports that TcdB increases spore adherence and internalization into intestinal tissue, the low Bezlotoxumab concentration used failed to fully protect against TcdB-mediated damage. These findings emphasize the need for further research to optimize therapeutic strategies for preventing CDI recurrence. The study also underscores the importance of dosage considerations in translating preclinical findings to clinical applications.

Based on our findings our proposed model (**Fig. 8**) illustrates the dual impact of *C. difficile* toxins and Bezlotoxumab treatment on intestinal epithelial cells. In untreated CDI, TcdA and TcdB intoxication leads to increased Acc Fn, Vn, and their associated integrins α_5_ and α_v_, ultimately resulting in enhanced spore adherence and internalization, alongside epithelial cell apoptosis. When Bezlotoxumab is administered, it partially neutralizes TcdB, reducing epithelial damage and spore adherence/internalization. However, the treatment still results in increased total Vn while decreasing total Fn, suggesting a complex modulation of these extracellular matrix proteins. This model supports our findings that low-dose Bezlotoxumab provides partial protection against TcdB-mediated damage while highlighting the intricate relationship between toxin activity and spore-host interactions.

**Fig. 8.**
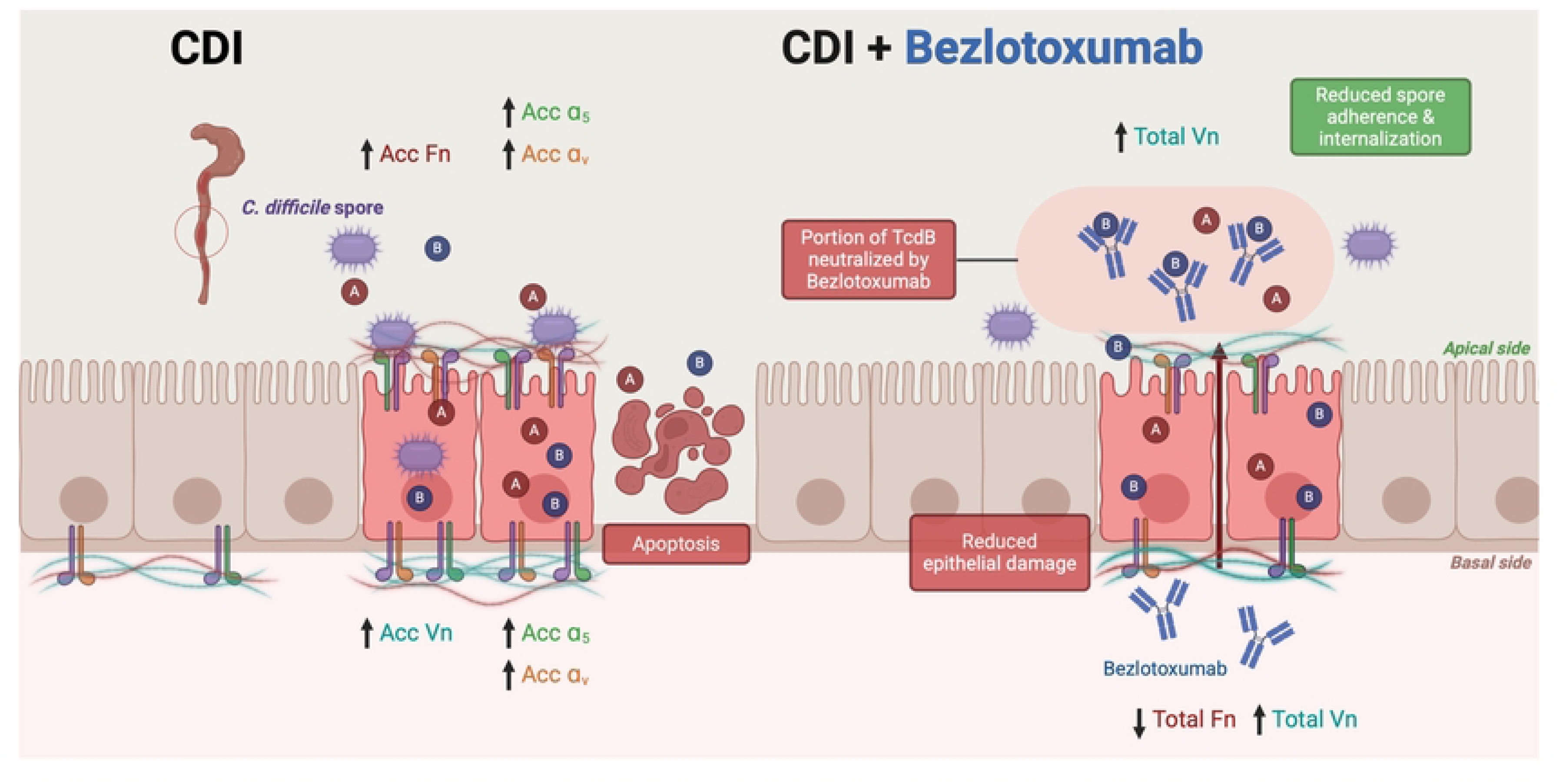
Model of *C. difficile* toxin-mediated spore interactions and Bezlotoxumab effects on intestinal epithelial cells. Left panel (CDI): During C. difficile infection, toxins A and B (TcdA/TcdB) cause increased accessible (Acc) fibronectin (Fn), vitronectin (Vn), and their associated integrins (α5 and αv) on both apical and basal surfaces of intestinal epithelial cells. This leads to enhanced spore adherence and internalization, ultimately resulting in epithelial cell apoptosis. Right panel (CDI + Bezlotoxumab): Treatment with Bezlotoxumab partially neutralizes TcdB, reducing epithelial damage and spore adherence/internalization. While total Vn levels increase, total Fn levels decrease. The presence of Bezlotoxumab antibodies results in reduced spore adherence and internalization compared to untreated conditions, demonstrating partial protection against TcdB- mediated damage. Arrows indicate increased (↑) or decreased (↓) levels of proteins. A and B represent TcdA and TcdB toxins, respectively. The apical and basal sides of the epithelium are indicated.

## Materials and Methods

### Data reporting

Statistical methods were not employed for predefining the sample size. In the case of animal experiments, mice were assigned to various groups through random allocation. The researchers did not employ blinding during the conduct of the animal experiments to prevent cross-contamination among animals from distinct treatment groups.

### Animals used in this study

Male or female C57BL/6 mice, aged 6-8 weeks, were sourced from the breeding colony at the Departamento de Ciencias Biológicas of the Universidad Andrés Bello, which was established using animals acquired from Jackson Laboratories. The mice were individually housed in cages maintained under uniform conditions, with autoclaved water, bedding, and cages. A 12-hour light-dark cycle was maintained, and the mice were kept at a temperature of 20-24°C with a humidity range of 40-60%. All experimental procedures strictly adhered to and were formally approved by the Institutional Animal Ethics Committee of the Universidad Andrés Bello (Protocol #015/2019) and the Animal Ethics Committee of the Facultad de Ciencias Biológicas of the Universidad Andrés Bello (Protocol #015/2019).

### Recombinant *C. difficile* toxin purification

TcdA and TcdB of *C. difficile* were produced in *Bacillus megaterium* carrying the plasmid pHIS1622, which harbors the cloned genes *tcdA* or *tcdB* from *C. difficile* VPI 10463, as previously documented^78^.

Transformed *B. megaterium* cells were cultivated in Luria-Bertani medium (BD, USA), supplemented with 10 µg/mL tetracycline. An overnight culture (35 mL) was used to inoculate 1 L of medium, and the bacteria were cultured at 37°C with shaking at 220 rpm. To induce toxin expression, 0.5% D-xylose was added once the culture reached an optical density at 600 nm of 0.5. After 4 hours, cells were harvested and resuspended in 200 mL of binding buffer (20 mM Tris [pH 8.0], 100 mM NaCl for TcdA and 20 mM Tris [pH 8.0] or 500 mM NaCl for TcdB), supplemented with 0.16 µg/mL DNase, 10 mg/mL lysozyme, and protease inhibitors (P8849; Sigma). Cell lysis was achieved using an Emulsiflex homogenizer, and lysates were centrifuged at 48,000 × g for 30 minutes. The toxins were purified from the supernatant through Ni-affinity, anion exchange, and size exclusion chromatography. The purified toxins were eluted and stored in 20 mM HEPES (pH 7.0), 50 mM NaCl.

### Purification of *C. difficile* spores

Spore preparation was conducted following previously established procedures^79^. In summary, a 1:1,000 dilution of an overnight culture in BHIS was plated (100 µL) on 70:30 agar plates (6.3% weight/vol BD, USA; 0.35% weight/vol protease peptone BD, USA; 0.07% ammonium sulfate (NH4)2SO4 Merck USA; 0.106% weight/vol Tris base Omnipur, Germany; 1.11% weight/vol brain heart infusion extract BD, USA; 0.15% weight/vol yeast extract BD, USA; 1.5% weight/vol Bacto agar BD, USA). Next, 70:30 agar plates were incubated for 7 days in an anaerobic chamber (Bactron III-2, Shellab USA) at 37°C. Subsequently, colonies were scraped from the plates with ice-cold sterile Milli-Q water. The sporulated culture underwent five washes with ice-cold Milli- Q water in a micro-centrifuge at 18,400×g for 5 minutes each. Spores were purified by density using 45% weight/vol autoclaved Nycodenz (Axell USA) and centrifugation at 18,400×g for 40 minutes. The bacterial debris and non-spore residuals were removed, and the spore pellet was separated and washed five times at 18,400×g for 5 minutes with ice-cold sterile Milli-Q water to eliminate Nycodenz. Spore concentration was determined in the Neubauer chamber, adjusted to 5 × 10^9^ spores/mL, and stored at −80°C until use.

### Immunofluorescence of E-cadherin, Fn, Vn, a_5_, a_v_, and β_1_ in TcdA and TcdB intoxicated Caco-2 cells

Caco-2 cells, cultured on glass coverslips in 24-well plates for 8 days post-confluence, underwent two washes with DPBS before intoxication with 600 pM of TcdA and TcdB (161,83pg/mL and 184.83 pg/mL) in DMEM FBS-Free or DMEM alone (as a control) for 3, 6, or 8 hours. Following treatment, cells were rinsed with PBS, fixed with PBS-4% paraformaldehyde, and further washed with PBS.

For staining surface-accessible proteins, cells were incubated with primary antibodies (1:200 dilution) including mouse monoclonal antibodies against human fibronectin (sc8422, Santa Cruz Biotechnologies, USA), vitronectin (sc74484, Santa Cruz Biotechnologies, USA), integrin α_5_ (ab78614, Abcam, USA), integrin α_v_ (ab16821, Abcam, USA), and integrin β_1_ (MAB1959Z, Millipore, USA). Incubation was carried out either overnight at 4 °C or for 1 hour at room temperature, followed by secondary antibody treatment using donkey anti-mouse IgG Alexa Fluor 488 (ab150109, Abcam, USA) at a 1:400 dilution.

To detect total protein, samples were permeabilized with PBS-0.2% Triton X-100 for 10 minutes, blocked with PBS-1% BSA, and subjected to the same primary antibody as used for surface staining. After washing, cells were incubated with donkey anti-mouse IgG Alexa Fluor 568 (ab175700, Abcam, USA). Finally, samples were mounted using Dako Fluorescent Mounting Medium (Dako) and visualized using confocal microscopy.

### Binding of Fn and Vn released from intoxicated cells to *C. difficile* spores

Differentiated Caco-2 cells were exposed to 600 pM of TcdA and TcdB for 8 hours at 37 °C in DMEM FBS-free. As a control, cells were incubated in DMEM FBS-free without toxins. Subsequently, the supernatant was meticulously purified of cellular debris through two centrifugation cycles: first at 760×g for 5 minutes, followed by collection of the supernatant, and then at 18,400×g for 10 minutes. This final supernatant was utilized in the subsequent experiment.

Next, 4 × 10^7^ *C. difficile* R20291 spores were incubated for 1 hour at 37 °C with 100 µL of the obtained supernatant from TcdA and TcdB-intoxicated cells or from cells incubated in DMEM FBS-free or fresh DMEM as a control. Unbound molecules were removed by washing the spores three times through cycles of centrifugation at 18,400×g for 5 minutes and resuspension in PBS.

The spores were then fixed on coverslips coated with poly-L-lysine, treated with PBS-4% PFA for 15 minutes at room temperature, and blocked with PBS-1% BSA for 1 hour at room temperature. Subsequently, coverslips were incubated with 1:200 dilution of mouse monoclonal anti-Fn (sc-8422, Santa Cruz Biotechnologies, USA) or mouse monoclonal anti-Vn (sc-74484, Santa Cruz Biotechnologies, USA) for 1 hour at room temperature. Following this, coverslips were incubated with 1:400 chicken anti-mouse IgG conjugated to Alexa 488 (A21200 ThermoFisher, USA) for 1 hour at room temperature. Finally, cells were mounted with Dako Fluorescent Mounting Medium (Dako). The samples were observed on an Olympus BX53 fluorescence microscope with UPLFLN 100× oil objective (numerical aperture 1.30), and images were captured using the microscope camera for fluorescence imaging, Qimaging R6 Retiga.

Quantitative analysis of Fn and Vn binding to *C. difficile* spores was conducted using ImageJ (NIH, USA), as previously described (see references Mora Uribe 2016 and Castro Cordova 2020). Relative Fn and Vn spore fluorescence intensity was measured by outlining around 600 spores per experimental condition, along with several adjacent background readings. The data presented represent [spore (Fl. int / area) – background (Fl. int / area)]. Fluorescence intensity profiles were generated from the microscopy images using the 3D Surface Plotter plug-in of ImageJ.

### Association of *C. difficile* spore with cellular Fibronectin or Vitronectin in differentiated Caco-2 cells intoxicated with TcdA and TcdB

To assess the interaction between *C. difficile* spores and cellular fibronectin or vitronectin during exposure to *C. difficile* toxins, differentiated Caco-2 cells were treated with 600 pM of TcdA and TcdB for 3, 6, or 8 hours at 37°C in DMEM FBS-free. As a control, cells were incubated in DMEM FBS-free. Subsequently, intoxicated cells were washed twice with PBS and then exposed to *C. difficile* spores for 3 hours at 37°C. Unbound spores were removed by three washes with PBS, and the cells were fixed with 4% PFA, followed by three additional washes with PBS.

In non-permeabilized monolayers, extracellular spores were labeled with 1:1,000 chicken IgY anti-*C. difficile* spore^80^ in PBS–1% BSA for 1 hour at room temperature. After two washes, cells were incubated with 1:400 goat anti-chicken IgY conjugated with Alexa Fluor 488 (ab150173, Abcam, USA) in PBS–1% BSA for 1 hour at room temperature. Following two washes, cells were stained with mouse monoclonal antibodies against human fibronectin (sc8422, Santa Cruz Biotechnologies, USA) or vitronectin (sc74484, Santa Cruz Biotechnologies, USA). Subsequently, samples were washed twice with PBS, and fibronectin or vitronectin were detected with 1:400 donkey anti-mouse IgG Alexa Fluor 568 (ab175700, Abcam, USA) for 1 hour at room temperature. After three washes with PBS and one with sterile distilled water, samples were dried at room temperature for 30 minutes, and coverslips were mounted using Dako Fluorescence Mounting Medium (Dako, Denmark) and sealed with nail polish. Samples were observed using confocal microscopy, as described below.

To determine the association of *C. difficile* spores with fibronectin and vitronectin released from differentiated Caco-2 cells intoxicated with TcdA and TcdB, cells were intoxicated for 8 hours in DMEM FBS-free. As a control, cells were treated with DMEM FBS-free. The supernatant was collected, carefully centrifuged at 760×g for 5 minutes, and then centrifuged at 18,400×g for 10 minutes. The new supernatant was collected, and 4 × 10^7^ *C. difficile* R20291 spores were incubated with 100 μL of the supernatant from intoxicated cells, healthy cells, or fresh DMEM for 1 hour at 37°C. Unbound molecules were washed off three times by cycles of centrifugation at 18,400×g for 5 minutes and resuspension in PBS. Spores were fixed on coverslips coated with poly-L-lysine, treated with PBS-4% PFA for 15 minutes at room temperature, and blocked with PBS-1% BSA for 1 hour at room temperature. Afterward, *C. difficile* spores and fibronectin or vitronectin were immunodetected as described above. Samples were observed on an Olympus BX53 fluorescence microscope with UPLFLN 100× oil objective (numerical aperture 1.30), and images were captured using the microscope camera for fluorescence imaging, Qimaging R6 Retiga. Quantitative analysis of Fn and Vn binding to *C. difficile* spores was conducted using ImageJ (NIH, USA), as previously published (Mora-Uribe et al., 2016). To measure the relative Fn and Vn spore fluorescence intensity, an outline was drawn around 600 spores per experimental condition and several adjacent background readings. The data shown correspond to [spore (Fl. Int / area) – background (Fl. int / area)]. Fluorescence intensity profiles were generated from the microscopy images using the 3D Surface Plotter plug-in of ImageJ.

Similarly, to investigate whether the association of Fn or Vn with *C. difficile* spores is time-dependent, differentiated Caco-2 cells were infected with *C. difficile* spores for 0, 15, 30, 60, 120, and 180 minutes at 37°C in DMEM FBS-free. Subsequently, cells were washed three times with PBS and fixed with 4% PFA. Immunostaining of spores and Fn or Vn was performed as described above. Samples were visualized in confocal microscopy, and the fluorescence intensity was quantified for each spore in the channels of anti-spore and anti-Fn or anti-Vn.

### Ileal loop assay

C57BL/6 mice were initially anesthetized in an isoflurane chamber (RWD USA) using 4% vol/vol isoflurane (Baxter USA) and maintained at 2% vol/vol during surgery administered by air. The established intestinal loop model was implemented as per previously referenced protocol^36, 81, 82^. Briefly, a midline laparotomy was performed, involving a 1-cm incision in the abdomen. The ileum and proximal colon (1.5 cm each, positioned 1.0 – 1.5 cm from the cecum as a reference) were ligated using silk surgical suture.

To assess the impact of TcdB on the cellular distribution of Fn and Vn in the ileum mucosa, C57BL/6 mice underwent ileum ligation and were injected with varying doses (0.1, 0.5, 1, or 5µg) of TcdB in 0.9% NaCl (saline), with a saline-only control (n = 3 for each treatment). For investigating whether TcdB influences the adherence of *C. difficile* spores in the ileum mucosa, ligated loops were injected with 5 × 10^8^ *C. difficile* R20291 spores and 1µg or 5µg of TcdB for 5 hours in saline (or saline alone as control) (n = 5 each treatment).

To explore the potential protective role of Bezlotoxumab in spore persistence in TcdB-intoxicated ileum, C57BL/6 mice were intraperitoneally injected with 5mg/kg of Bezlotoxumab (n = 10) 24 hours before surgery, with saline as a control. Subsequently, Bezlotoxumab-treated mice were divided into two groups: one injected with 5 × 10^8^ *C. difficile* R20291 spores (n = 5), and the other with both spores and 5µg of TcdB (n = 5). For animals injected with saline, loops were injected with 5 × 10^8^ *C. difficile* R20291 spores and 5µg of TcdB (n = 5). Following the procedures, the intestine was returned to the abdomen, the incision was closed, and the animals were allowed to regain consciousness. Mice were kept for 5 hours before euthanasia. The ligated loops were carefully removed and washed in PBS before subsequent immunostaining, as detailed below.

### Immunostaining of ileal and colonic loops

Initially, we longitudinally cut the extracted and washed intestinal tissues from the ileal and colonic loops. Subsequently, these tissues underwent a triple immersion in PBS at room temperature (RT) for thorough washing. For enhanced visualization, the tissues were flattened at RT. This involved fixing the tissues flat over a filter paper saturated with a solution containing 30% sucrose (Winkler, Chile) in PBS–4% paraformaldehyde (Merck, USA) for a minimum of 15 minutes. Following this, the tissues were transferred to a microcentrifuge tube containing the same fixing solution and were left to incubate at 4 °C overnight.

Given that fixing mucus with cross-linking agents like paraformaldehyde can lead to the collapse and shrinkage of the mucus layer in the colon^83^, it’s noteworthy that we did not observe a mucus layer in our ileal and colonic loops. In preparation for immunostaining, the intestinal and colonic tissues were subsequently cut into approximately ∼5 × 5 mm fragments.

In the TcdB-intoxicated ileum, for immunostaining of both luminally accessible and total fibronectin or vitronectin, ileum fragments underwent an incubation step with a primary antibody at a dilution of 1:150 rabbit pAb anti-fibronectin (sc-9068, Santa Cruz Biotechnology, USA) or rabbit pAb anti-vitronectin (sc-15332, Santa Cruz Biotechnology, USA) in PBS-3% BSA overnight at 4 °C. Subsequent to washing, the fragments were incubated with a secondary antibody at a dilution of 1:400 donkey anti-rabbit IgG Alexa 568 (ab175692, Abcam, USA) for 3 hours at RT. To stain total protein, the tissues were permeabilized through a 2-hour incubation with PBS- 0.2% Triton X-100 at RT and then blocked with 3% BSA for 3 hours at RT. Following this, the tissues were incubated overnight at 4 °C with the same primary antibody used earlier (1:150). F- actin was stained using a 1:150 dilution of fluorescently labeled phalloidin Alexa Fluor 647 (A22287, ThermoFisher, USA). On the subsequent day, the tissues were washed and incubated with a mixture of 1:400 donkey anti-rabbit IgG Alexa 488 (ab150061, Abcam, USA) and Hoechst at a 1:1,000 dilution for 3 hours at room temperature.

For the quantification of *C. difficile* spore adherence and internalization in both colonic and ileum mucosa, tissues underwent permeabilization through incubation with PBS–0.2% Triton X-100 (Merck, USA) and were subsequently blocked with PBS–3% BSA (Sigma–Aldrich, USA) for 3 hours at room temperature (RT). The same buffer was utilized for the subsequent antibody incubation. The tissue was then exposed to a primary polyclonal antibody, specifically a 1:1,000 dilution of anti-*C. difficile* spore IgY batch 7246 antibodies (Aveslab USA) in PBS–3% BSA. Notably, these antibodies do not react with epitopes of vegetative cells or murine microbiota^80^. Concurrently, the tissue was also incubated with a 1:50 dilution of phalloidin Alexa-Fluor 568 (#ab176753 Abcam, USA) in PBS–3% BSA overnight at 4 °C to stain the actin cytoskeleton.

Following a PBS wash, the samples were incubated with a secondary antibody, specifically a 1:400 dilution of goat anti-chicken IgY Alexa-Fluor 488 (#ab150173 Abcam USA) in PBS–3% BSA at RT. After three PBS washes, cellular nuclei were stained with a 1:1,000 dilution of Hoechst (ThermoFisher, USA) for 15 minutes at RT. Subsequently, the immune-stained tissues were mounted with the luminal side facing up, following the protocol previously described^81^.

### Confocal and Epifluorescence analysis of immunostained tissued or intestinal epithelial cells

For confocal imaging, the Leica SP8 system was employed, utilizing an HPL APO CS2 40× oil objective with a numerical aperture of 1.30. Detection of signals was facilitated through three PMT spectral detectors: PMT1 (410-483) for DAPI, PMT2 (505-550) for Alexa-Fluor 488, and PMT3 (587-726) for Alexa-Fluor 555. Emitted fluorescence was separated using dichroic mirrors DD488/552.

To quantify the fluorescence intensity of fibronectin, vitronectin, α5, αv, and β1 images (1,024×1,024 pixels) were acquired with a 0.5-μm Z step. Fluorescence intensity measurements were conducted for channels representing both accessible and total protein. The z-stack analysis involved an equal number of slides. To assess fluorescence intensity in the basal and apical halves, z-stacks were divided into halves with an equal number of slides, and the fluorescence intensity was then summed. Six microscopy fields were analyzed for each experimental condition.

For the evaluation of luminally accessible fibronectin and vitronectin in the ileum mucosa, images of 1,024×1,024 pixels were captured with a 2-μm Z step size, followed by filtering with Gaussian Blur 3D (sigma x: 0.6; y: 0.6; z: 0.6). Three-dimensional reconstructions of the ileum mucosa were performed using ImageJ software (NIH, USA). Villi were visualized through Hoechst and phalloidin signals. Each animal contributed two pictures (n = 3 independent mice), and the total intestinal epithelium area analyzed for each mouse was 0.5 mm². Representative 3D projections were generated using the 3D projection plug-in of ImageJ.

## Acknowledgments

Thanks to members of the Paredes-Sabja laboratory for their helpful comments during the preparation of this manuscript. This project was supported by Merck Sharp & Dohme LLC, a subsidiary of Merck & Co., Inc., Rahway, NJ, USA, and awards 1R01AI177842 from the National Institute of Allergy and Infectious Diseases to D.P-S. The content is solely the responsibility of the authors and does not necessarily represent the official views of the NIAID. The funders had no role in study design, data collection and interpretation, or the decision to submit the work for publication.

---

**Fig. S1.**
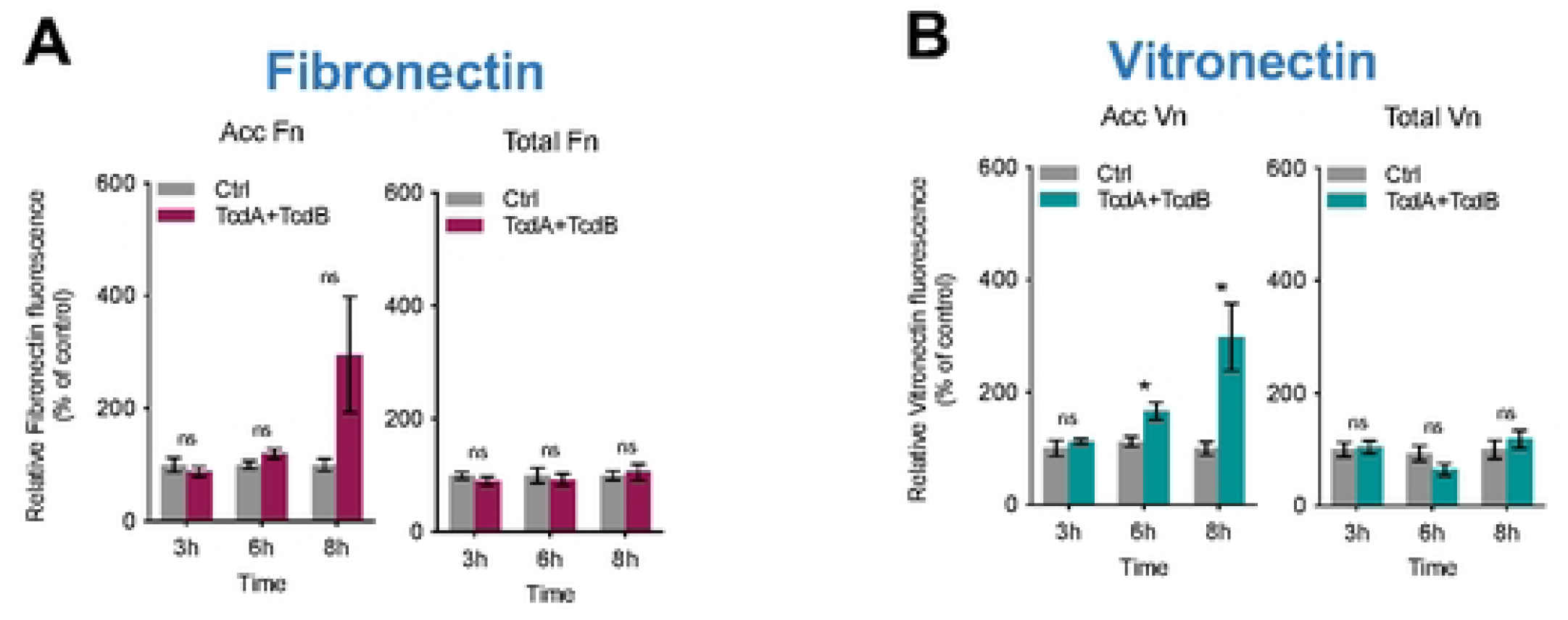
Effect of TcdA/TcdB intoxication of intestinal epithelial Caco-2 cells in levels of Fibronectin and Vitronectin. Differentiated Caco-2 cells intoxicated with TcdA and TcdB for 3, 6, or 8h in DMEM FBS-free. As a control, cells were treated with DMEM FBS-free. Relative fluorescence intensity measured as the sum of raw intensity density/area for each z-step of accessible **a** (acc) and total Fn, its abundance in the cell; in the same way, the relative fluorescence intensity of **b** (acc) and total Vn, its abundance in the cell. Controls were set at 100%. Error bars indicate the mean ± SEM from at least 9 fields (*n* = 3). Statistical analysis was performed by Two-Way ANOVA post-Bonferroni; ns, *p* > 0.05; * *p* < 0.05.

**Fig. S2.**
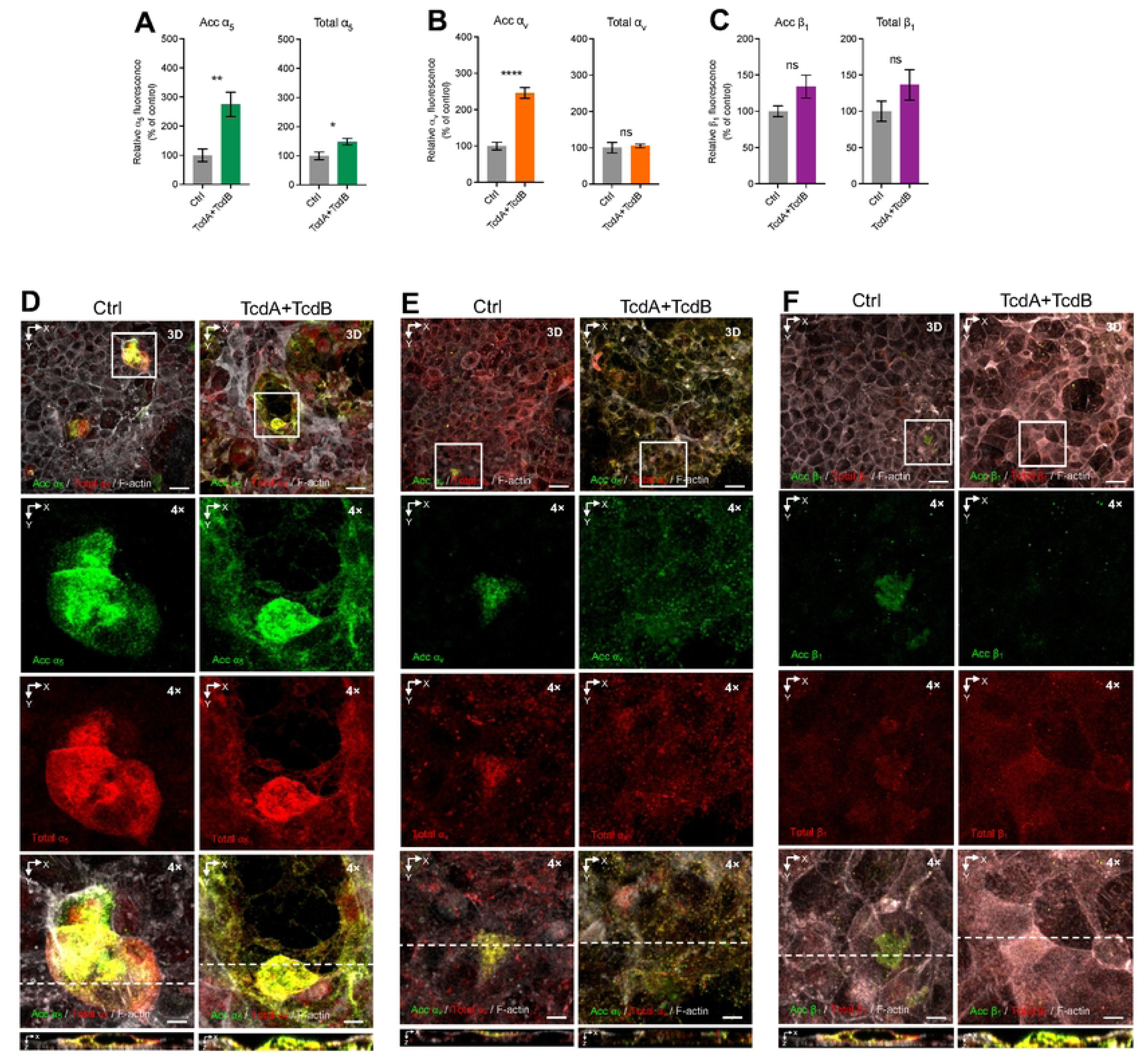
TcdA and TcdB increase accessible ⍺_5_ and ⍺_V_ but no β_1_ integrins in intestinal epithelial cells. Differentiated Caco-2 cells intoxicated with 600pM of TcdA and TcdB for 8h in DMEM FBS-free. As a control, cells were treated with DMEM FBS-free. Unpermeabilized cells were stained for accessible integrin (green), permeabilized, and stained total integrin (red) and F-actin (grey). Cells were immunostained for ⍺_v_ integrin; Relative fluorescence intensity measured as the sum of raw intensity density/area for each z-step of **a,** acc ⍺_5_ and total ⍺_5_ in cells, **b**, acc ⍺_V_ and total ⍺_V_ in cells,**c,** acc β_1_ and total β_1_ in cells. **d, e, f**, Representative confocal microscopy images 3D projection of control cells (left) and intoxicated cells for 8h (right); below 4x magnified slides (XY), and the orthogonal view (XZ). Controls were set 100%. Error bars indicate the mean ± S.E.M from at least 9 fields (*n* = 3). Statistical analysis was performed by unpaired Student’s *t* test, ns indicates non-significant differences; ** *p* < 0.01; **** *p* < <0.0001. Bars, top panels 20 µm; bottom panels 5µm.

**Fig. S3.**
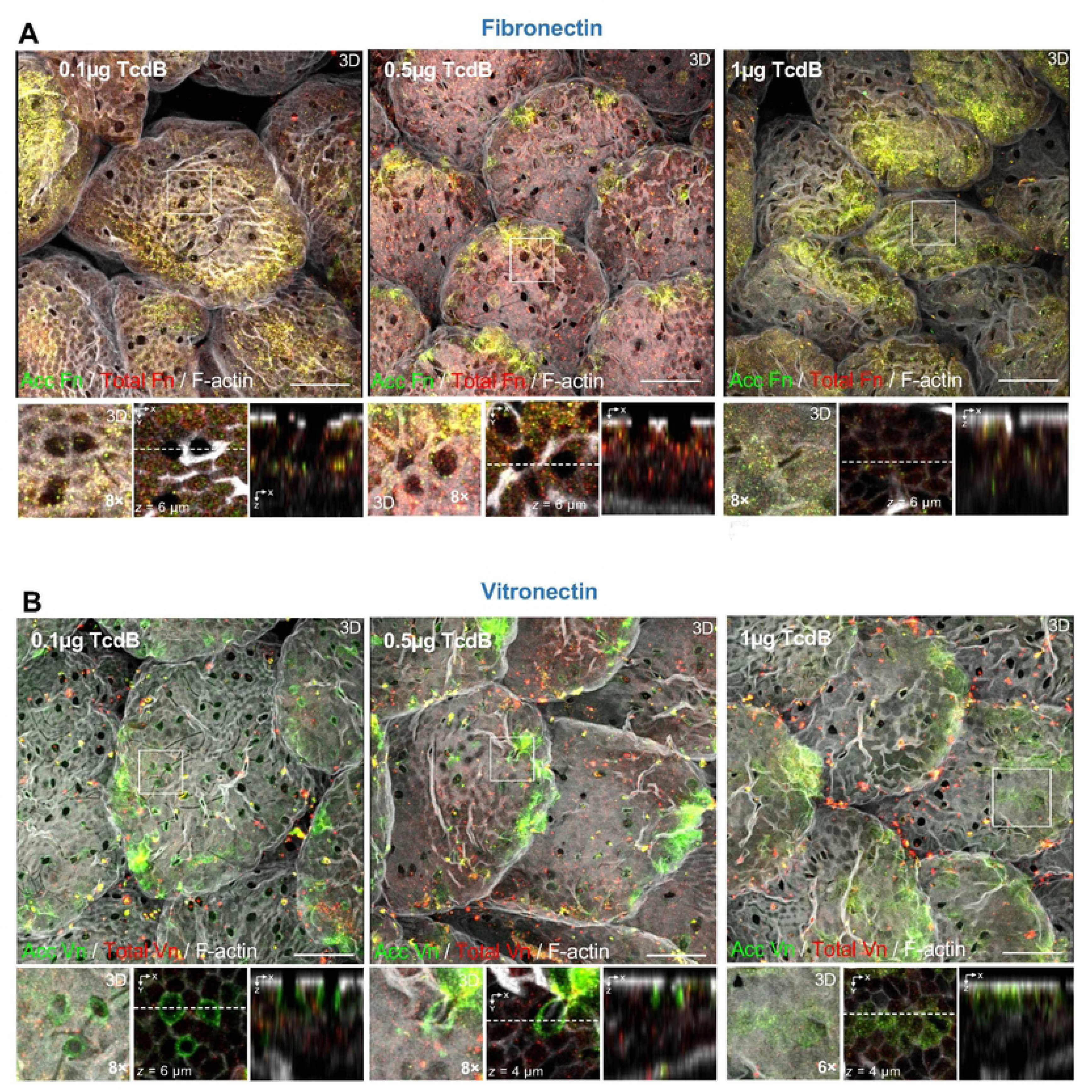
Effect of TcdB-intoxicated ileal loop in accessible and total levels of Fibronectin and Vitronectin. Ileal ligated loops were intoxicated for 5 h with 0.1, 0.5, 1, or 5µg of TcdB or saline as control. Then loops were removed, washed, fixed, and subjected to immunofluorescence. Unpermeabilized tissues were stained for accessible Fibronectin or Vitronectin (acc Fn or acc Vn; green), and then permeabilized and stained total Fibronectin or Vitronectin (total Fn or total Vn; red) and F-actin (grey). **a**-**b**, Representative confocal microscopy images 3D projection of intoxicated loops for 5 h with 0.1µg (left), intoxicated loops for 5 h with 0.5µg (middle), and intoxicated loops for 5 h with 0.1µg (right) immunostained for accessible and total Fn or Vn; right bottom, a magnified 3D projection, next to a magnified z-stack (XY), and then magnified orthogonal view (XZ). Scale bar 20 µm.

## References

1. Lessa FC, Mu Y, Bamberg WM, Beldavs ZG, Dumyati GK, Dunn JR, Farley MM, Holzbauer SM, Meek JI, Phipps EC, Wilson LE, Winston LG, Cohen JA, Limbago BM, Fridkin SK, Gerding DN, McDonald LC. Burden of *Clostridium difficile* infection in the United States. N Engl J Med. 2015;372(9):825–34. Epub 2015/02/26. doi: 10.1056/NEJMoa1408913. PubMed PMID: 25714160.

2. McDonald LC, Gerding DN, Johnson S, Bakken JS, Carroll KC, Coffin SE, Dubberke ER, Garey KW, Gould CV, Kelly C, Loo V, Shaklee Sammons J, Sandora TJ, Wilcox MH. Clinical Practice Guidelines for Clostridium difficile Infection in Adults and Children: 2017 Update by the Infectious Diseases Society of America (IDSA) and Society for Healthcare Epidemiology of America (SHEA). Clin Infect Dis. 2018;66(7):987–94. doi: 10.1093/cid/ciy149. PubMed PMID: 29562266.

3. Kelly CP. Can we identify patients at high risk of recurrent Clostridium difficile infection? Clin Microbiol Infect. 2012;18 Suppl 6:21–7. doi: 10.1111/1469-0691.12046. PubMed PMID: 23121551.

4. Zhang S, Palazuelos-Munoz S, Balsells EM, Nair H, Chit A, Kyaw MH. Cost of hospital management of Clostridium difficile infection in United States-a meta-analysis and modelling study. BMC Infect Dis. 2016;16(1):447. Epub 20160825. doi: 10.1186/s12879-016-1786-6. PubMed PMID: 27562241; PMCID: PMC5000548.

5. Guery B, Galperine T, Barbut F. *Clostridioides difficile*: diagnosis and treatments. BMJ. 2019;366:l4609. Epub 2019/08/20. doi: 10.1136/bmj.l4609. PubMed PMID: 31431428.

6. Schnizlein MK, Young VB. Capturing the environment of the Clostridioides difficile infection cycle. Nat Rev Gastroenterol Hepatol. 2022;19(8):508–20. Epub 20220425. doi: 10.1038/s41575-022-00610-0. PubMed PMID: 35468953.

7. Aktories K, Schwan C, Jank T. *Clostridium difficile* toxin biology. Annu Rev Microbiol. 2017;71:281–307. Epub 2017/06/29. doi: 10.1146/annurev-micro-090816-093458. PubMed PMID: 28657883.

8. Smits WK, Lyras D, Lacy DB, Wilcox MH, Kuijper EJ. Clostridium difficile infection. Nat Rev Dis Primers. 2016;2:16020. Epub 20160407. doi: 10.1038/nrdp.2016.20. PubMed PMID: 27158839; PMCID: PMC5453186.

9. Paredes-Sabja D, Shen A, Sorg JA. *Clostridium difficile* spore biology: sporulation, germination, and spore structural proteins. Trends Microbiol. 2014;22(7):406–16. Epub 2014/05/13. doi: 10.1016/j.tim.2014.04.003. PubMed PMID: 24814671; PMCID: PMC4098856.

10. Martens EC, Neumann M, Desai MS. Interactions of commensal and pathogenic microorganisms with the intestinal mucosal barrier. Nat Rev Microbiol. 2018;16(8):457–70. doi: 10.1038/s41579-018-0036-x. PubMed PMID: 29904082.

11. Singh B, Su YC, Riesbeck K. Vitronectin in bacterial pathogenesis: a host protein used in complement escape and cellular invasion. Mol Microbiol. 2010;78(3):545–60. Epub 2010/09/03. doi: 10.1111/j.1365-2958.2010.07373.x. PubMed PMID: 20807208.

12. Henderson B, Nair S, Pallas J, Williams MA. Fibronectin: a multidomain host adhesin targeted by bacterial fibronectin-binding proteins. FEMS Microbiol Rev. 2011;35(1):147–200. Epub 2010/08/11. doi: 10.1111/j.1574-6976.2010.00243.x. PubMed PMID: 20695902.

13. Quaroni A, Isselbacher KJ, Ruoslahti E. Fibronectin synthesis by epithelial crypt cells of rat small intestine. Proc Natl Acad Sci U S A. 1978;75(11):5548–52. doi: 10.1073/pnas.75.11.5548. PubMed PMID: 103096; PMCID: PMC393003.

14. Claudepierre P, Allanore Y, Belec L, Larget-Piet B, Zardi L, Chevalier X. Increased Ed-B fibronectin plasma levels in spondyloarthropathies: comparison with rheumatoid arthritis patients and a healthy population. Rheumatology (Oxford). 1999;38(11):1099–103. doi: 10.1093/rheumatology/38.11.1099. PubMed PMID: 10556262.

15. Vachon PH, Simoneau A, Herring-Gillam FE, Beaulieu JF. Cellular fibronectin expression is down-regulated at the mRNA level in differentiating human intestinal epithelial cells. Exp Cell Res. 1995;216(1):30–4. doi: 10.1006/excr.1995.1004. PubMed PMID: 7813630.

16. Hayman EG, Pierschbacher MD, Ohgren Y, Ruoslahti E. Serum spreading factor (vitronectin) is present at the cell surface and in tissues. Proc Natl Acad Sci U S A. 1983;80(13):4003–7. doi: 10.1073/pnas.80.13.4003. PubMed PMID: 6191326; PMCID: PMC394188.

17. Pellegrini A, Pietrocola G. Recruitment of Vitronectin by Bacterial Pathogens: A Comprehensive Overview. Microorganisms. 2024;12(7). Epub 20240708. doi: 10.3390/microorganisms12071385. PubMed PMID: 39065153; PMCID: PMC11278874.

18. Schvartz I, Seger D, Shaltiel S. Vitronectin. Int J Biochem Cell Biol. 1999;31(5):539–44. doi: 10.1016/s1357-2725(99)00005-9. PubMed PMID: 10399314.

19. Podack ER, Preissner KT, Müller-Eberhard HJ. Inhibition of C9 polymerization within the SC5b-9 complex of complement by S-protein. Acta Pathol Microbiol Immunol Scand Suppl. 1984;284:89–96. PubMed PMID: 6587746.

20. Peters JH, Loredo GA, Chen G, Maunder R, Hahn TJ, Willits NH, Hynes RO. Plasma levels of fibronectin bearing the alternatively spliced EIIIB segment are increased after major trauma. J Lab Clin Med. 2003;141(6):401–10. doi: 10.1016/s0022-2143(03)00042-8. PubMed PMID: 12819638.

21. Castellanos M, Leira R, Serena J, Blanco M, Pedraza S, Castillo J, Dávalos A. Plasma cellular-fibronectin concentration predicts hemorrhagic transformation after thrombolytic therapy in acute ischemic stroke. Stroke. 2004;35(7):1671–6. Epub 20040527. doi: 10.1161/01.str.0000131656.47979.39. PubMed PMID: 15166391.

22. Kolachala VL, Bajaj R, Wang L, Yan Y, Ritzenthaler JD, Gewirtz AT, Roman J, Merlin D, Sitaraman SV. Epithelial-derived fibronectin expression, signaling, and function in intestinal inflammation. J Biol Chem. 2007;282(45):32965–73. Epub 2007/09/15. doi: 10.1074/jbc.M704388200. PubMed PMID: 17855340.

23. Giuffrida P, Pinzani M, Corazza GR, Di Sabatino A. Biomarkers of intestinal fibrosis - one step towards clinical trials for stricturing inflammatory bowel disease. United European Gastroenterol J. 2016;4(4):523–30. Epub 20160321. doi: 10.1177/2050640616640160. PubMed PMID: 27536362; PMCID: PMC4971797.

24. Pan W, Xiang L, Liang X, Du W, Zhao J, Zhang S, Zhou X, Geng L, Gong S, Xu W. Vitronectin Destroyed Intestinal Epithelial Cell Differentiation through Activation of PDE4- Mediated Ferroptosis in Inflammatory Bowel Disease. Mediators Inflamm. 2023;2023:6623329. Epub 20230719. doi: 10.1155/2023/6623329. PubMed PMID: 37501933; PMCID: PMC10371469.

25. Schwan C, Kruppke AS, Nolke T, Schumacher L, Koch-Nolte F, Kudryashev M, Stahlberg H, Aktories K. *Clostridium difficile* toxin CDT hijacks microtubule organization and reroutes vesicle traffic to increase pathogen adherence. Proc Natl Acad Sci U S A. 2014;111(6):2313–8. Epub 2014/01/29. doi: 10.1073/pnas.1311589111. PubMed PMID: 24469807; PMCID: PMC3926047.

26. Schwan C, Stecher B, Tzivelekidis T, van Ham M, Rohde M, Hardt WD, Wehland J, Aktories K. Clostridium difficile toxin CDT induces formation of microtubule-based protrusions and increases adherence of bacteria. PLoS Pathog. 2009;5(10):e1000626. Epub 20091016. doi: 10.1371/journal.ppat.1000626. PubMed PMID: 19834554; PMCID: PMC2757728.

27. Rupnik M, Wilcox MH, Gerding DN. Clostridium difficile infection: new developments in epidemiology and pathogenesis. Nat Rev Microbiol. 2009;7(7):526–36. doi: 10.1038/nrmicro2164. PubMed PMID: 19528959.

28. Villafuerte Gálvez JA, Kelly CP. Bezlotoxumab: anti-toxin B monoclonal antibody to prevent recurrence of Clostridium difficile infection. Expert Rev Gastroenterol Hepatol. 2017;11(7):611–22. Epub 20170626. doi: 10.1080/17474124.2017.1344551. PubMed PMID:28636484.

29. Zhang Z, Chen X, Hernandez LD, Lipari P, Flattery A, Chen SC, Kramer S, Polishook JD, Racine F, Cape H, Kelly CP, Therien AG. Toxin-mediated paracellular transport of antitoxin antibodies facilitates protection against Clostridium difficile infection. Infect Immun. 2015;83(1):405–16. Epub 20141110. doi: 10.1128/iai.02550-14. PubMed PMID: 25385797; PMCID: PMC4288887.

30. Kyne L, Warny M, Qamar A, Kelly CP. Association between antibody response to toxin A and protection against recurrent Clostridium difficile diarrhoea. Lancet. 2001;357(9251):189-93. doi: 10.1016/s0140-6736(00)03592-3. PubMed PMID: 11213096.

31. Kyne L, Warny M, Qamar A, Kelly CP. Asymptomatic carriage of Clostridium difficile and serum levels of IgG antibody against toxin A. N Engl J Med. 2000;342(6):390–7. doi: 10.1056/nejm200002103420604. PubMed PMID: 10666429.

32. Lowy I, Molrine DC, Leav BA, Blair BM, Baxter R, Gerding DN, Nichol G, Thomas WD, Jr., Leney M, Sloan S, Hay CA, Ambrosino DM. Treatment with monoclonal antibodies against Clostridium difficile toxins. N Engl J Med. 2010;362(3):197–205. doi: 10.1056/NEJMoa0907635. PubMed PMID: 20089970.

33. Castro-Córdova P, Otto-Medina M, Montes-Bravo N, Brito-Silva C, Lacy DB, Paredes- Sabja D. Redistribution of the Novel Clostridioides difficile Spore Adherence Receptor E- Cadherin by TcdA and TcdB Increases Spore Binding to Adherens Junctions. Infect Immun. 2023;91(1):e0047622. Epub 20221130. doi: 10.1128/iai.00476-22. PubMed PMID: 36448839; PMCID: PMC9872679.

34. Janvilisri T, Scaria J, Chang YF. Transcriptional profiling of *Clostridium difficile* and Caco-2 cells during infection. J Infect Dis. 2010;202(2):282–90. Epub 2010/06/05. doi: 10.1086/653484. PubMed PMID: 20521945; PMCID: PMC2891111.

35. D’Auria KM, Kolling GL, Donato GM, Warren CA, Gray MC, Hewlett EL, Papin JA. In vivo physiological and transcriptional profiling reveals host responses to *Clostridium difficile* toxin A and toxin B. Infect Immun. 2013;81(10):3814–24. Epub 2013/07/31. doi: 10.1128/IAI.00869-13. PubMed PMID: 23897615; PMCID: PMC3811747.

36. Castro-Córdova P, Mora-Uribe P, Reyes-Ramírez R, Cofré-Araneda G, Orozco-Aguilar J, Brito-Silva C, Mendoza-León MJ, Kuehne SA, Minton NP, Pizarro-Guajardo M, Paredes-Sabja D. Entry of spores into intestinal epithelial cells contributes to recurrence of *Clostridioides difficile* infection. Nat Commun. 2021;12(1):1140. Epub 2021/02/18. doi: 10.1038/s41467-021-21355-5. PubMed PMID: 33602902.

37. Niederlechner S, Klawitter J, Baird C, Kallweit AR, Christians U, Wischmeyer PE. Fibronectin-integrin signaling is required for L-glutamine’s protection against gut injury. PLoS One. 2012;7(11):e50185. Epub 20121120. doi: 10.1371/journal.pone.0050185. PubMed PMID: 23185570; PMCID: PMC3502344.

38. Abdullayeva G, Liu H, Liu TC, Simmons A, Novelli M, Huseynova I, Lastun VL, Bodmer W. Goblet cell differentiation subgroups in colorectal cancer. Proc Natl Acad Sci U S A. 2024;121(43):e2414213121. Epub 20241014. doi: 10.1073/pnas.2414213121. PubMed PMID:

39. 39401352; PMCID: PMC11513979.

39. Mora-Uribe P, Miranda-Cardenas C, Castro-Cordova P, Gil F, Calderon I, Fuentes JA, Rodas PI, Banawas S, Sarker MR, Paredes-Sabja D. Characterization of the Adherence of *Clostridium difficile* Spores: The Integrity of the Outermost Layer Affects Adherence Properties of Spores of the Epidemic Strain R20291 to Components of the Intestinal Mucosa. Front Cell Infect Microbiol. 2016;6:99. Epub 2016/10/08. doi: 10.3389/fcimb.2016.00099. PubMed PMID: 27713865; PMCID: PMC5031699.

40. Zeng Z, Zhao H, Dorr MB, Shen J, Wilcox MH, Poxton IR, Guris D, Li J, Shaw PM. Bezlotoxumab for prevention of Clostridium difficile infection recurrence: Distinguishing relapse from reinfection with whole genome sequencing. Anaerobe. 2020;61:102137. Epub 2019/12/14. doi: 10.1016/j.anaerobe.2019.102137. PubMed PMID: 31846705.

41. Wilcox MH, Gerding DN, Poxton IR, Kelly C, Nathan R, Birch T, Cornely OA, Rahav G, Bouza E, Lee C, Jenkin G, Jensen W, Kim YS, Yoshida J, Gabryelski L, Pedley A, Eves K, Tipping R, Guris D, Kartsonis N, Dorr MB, Modify I, Investigators MI. Bezlotoxumab for Prevention of Recurrent *Clostridium difficile* Infection. N Engl J Med. 2017;376(4):305–17. Epub 2017/01/26. doi: 10.1056/NEJMoa1602615. PubMed PMID: 28121498.

42. Denny JE, Alam MZ, Mdluli NV, Maslanka JR, Lieberman LA, Abt MC. Monoclonal antibody-mediated neutralization of Clostridioides difficile toxin does not diminish induction of the protective innate immune response to infection. Anaerobe. 2024;88:102859. Epub 20240501. doi: 10.1016/j.anaerobe.2024.102859. PubMed PMID: 38701911; PMCID: PMC11347114.

43. Deakin LJ, Clare S, Fagan RP, Dawson LF, Pickard DJ, West MR, Wren BW, Fairweather NF, Dougan G, Lawley TD. The Clostridium difficile spo0A gene is a persistence and transmission factor. Infect Immun. 2012;80(8):2704–11. Epub 20120521. doi: 10.1128/iai.00147-12. PubMed PMID: 22615253; PMCID: PMC3434595.

44. Nikitas G, Deschamps C, Disson O, Niault T, Cossart P, Lecuit M. Transcytosis of Listeria monocytogenes across the intestinal barrier upon specific targeting of goblet cell accessible E- cadherin. J Exp Med. 2011;208(11):2263–77. Epub 20111003. doi: 10.1084/jem.20110560. PubMed PMID: 21967767; PMCID: PMC3201198.

45. Pentecost M, Kumaran J, Ghosh P, Amieva MR. Listeria monocytogenes internalin B activates junctional endocytosis to accelerate intestinal invasion. PLoS Pathog. 2010;6(5):e1000900. Epub 20100513. doi: 10.1371/journal.ppat.1000900. PubMed PMID: 20485518; PMCID: PMC2869327.

46. Pentecost M, Otto G, Theriot JA, Amieva MR. Listeria monocytogenes invades the epithelial junctions at sites of cell extrusion. PLoS Pathog. 2006;2(1):e3. Epub 20060127. doi: 10.1371/journal.ppat.0020003. PubMed PMID: 16446782; PMCID: PMC1354196.

47. Just I, Selzer J, Wilm M, von Eichel-Streiber C, Mann M, Aktories K. Glucosylation of Rho proteins by Clostridium difficile toxin B. Nature. 1995;375(6531):500-3. doi: 10.1038/375500a0. PubMed PMID: 7777059.

48. Chaves-Olarte E, Weidmann M, Eichel-Streiber C, Thelestam M. Toxins A and B from Clostridium difficile differ with respect to enzymatic potencies, cellular substrate specificities, and surface binding to cultured cells. J Clin Invest. 1997;100(7):1734–41. doi: 10.1172/jci119698. PubMed PMID: 9312171; PMCID: PMC508356.

49. Jefferson KK, Smith MF, Jr., Bobak DA. Roles of intracellular calcium and NF-kappa B in the Clostridium difficile toxin A-induced up-regulation and secretion of IL-8 from human monocytes. J Immunol. 1999;163(10):5183–91. PubMed PMID: 10553038.

50. Lee JY, Kim H, Cha MY, Park HG, Kim YJ, Kim IY, Kim JM. Clostridium difficile toxin A promotes dendritic cell maturation and chemokine CXCL2 expression through p38, IKK, and the NF-kappaB signaling pathway. J Mol Med (Berl). 2009;87(2):169–80. Epub 20081105. doi: 10.1007/s00109-008-0415-2. PubMed PMID: 18985311.

51. Xu H, Yang J, Gao W, Li L, Li P, Zhang L, Gong YN, Peng X, Xi JJ, Chen S, Wang F, Shao F. Innate immune sensing of bacterial modifications of Rho GTPases by the Pyrin inflammasome. Nature. 2014;513(7517):237-41. Epub 20140611. doi: 10.1038/nature13449. PubMed PMID: 24919149.

52. Jafari NV, Kuehne SA, Bryant CE, Elawad M, Wren BW, Minton NP, Allan E, Bajaj- Elliott M. Clostridium difficile modulates host innate immunity via toxin-independent and dependent mechanism(s). PLoS One. 2013;8(7):e69846. Epub 20130729. doi: 10.1371/journal.pone.0069846. PubMed PMID: 23922820; PMCID: PMC3726775.

53. Ng J, Hirota SA, Gross O, Li Y, Ulke-Lemee A, Potentier MS, Schenck LP, Vilaysane A, Seamone ME, Feng H, Armstrong GD, Tschopp J, Macdonald JA, Muruve DA, Beck PL. Clostridium difficile toxin-induced inflammation and intestinal injury are mediated by the inflammasome. Gastroenterology. 2010;139(2):542–52, 52.e1-3. Epub 20100413. doi: 10.1053/j.gastro.2010.04.005. PubMed PMID: 20398664.

54. Chumbler NM, Farrow MA, Lapierre LA, Franklin JL, Haslam DB, Goldenring JR, Lacy DB. Clostridium difficile Toxin B causes epithelial cell necrosis through an autoprocessing- independent mechanism. PLoS Pathog. 2012;8(12):e1003072. Epub 20121206. doi: 10.1371/journal.ppat.1003072. PubMed PMID: 23236283; PMCID: PMC3516567.

55. Qa’Dan M, Ramsey M, Daniel J, Spyres LM, Safiejko-Mroczka B, Ortiz-Leduc W, Ballard JD. Clostridium difficile toxin B activates dual caspase-dependent and caspase-independent apoptosis in intoxicated cells. Cell Microbiol. 2002;4(7):425–34. doi: 10.1046/j.1462-5822.2002.00201.x. PubMed PMID: 12102688.

56. Farrow MA, Chumbler NM, Lapierre LA, Franklin JL, Rutherford SA, Goldenring JR, Lacy DB. Clostridium difficile toxin B-induced necrosis is mediated by the host epithelial cell NADPH oxidase complex. Proc Natl Acad Sci U S A. 2013;110(46):18674–9. Epub 20131028. doi: 10.1073/pnas.1313658110. PubMed PMID: 24167244; PMCID: PMC3831945.

57. Wohlan K, Goy S, Olling A, Srivaratharajan S, Tatge H, Genth H, Gerhard R. Pyknotic cell death induced by Clostridium difficile TcdB: chromatin condensation and nuclear blister are induced independently of the glucosyltransferase activity. Cell Microbiol. 2014;16(11):1678–92. Epub 20140804. doi: 10.1111/cmi.12317. PubMed PMID: 24898616.

58. Frazer LC, Good M. Intestinal epithelium in early life. Mucosal Immunol. 2022;15(6):1181–7. Epub 20221115. doi: 10.1038/s41385-022-00579-8. PubMed PMID: 36380094; PMCID: PMC10329854.

59. Kordus SL, Thomas AK, Lacy DB. Clostridioides difficile toxins: mechanisms of action and antitoxin therapeutics. Nat Rev Microbiol. 2022;20(5):285–98. Epub 20211126. doi: 10.1038/s41579-021-00660-2. PubMed PMID: 34837014; PMCID: PMC9018519.

60. Mileto SJ, Jardé T, Childress KO, Jensen JL, Rogers AP, Kerr G, Hutton ML, Sheedlo MJ, Bloch SC, Shupe JA, Horvay K, Flores T, Engel R, Wilkins S, McMurrick PJ, Lacy DB, Abud HE, Lyras D. *Clostridioides difficile* infection damages colonic stem cells via TcdB, impairing epithelial repair and recovery from disease. Proc Natl Acad Sci U S A. 2020;117(14):8064–73. Epub 2020/03/20. doi: 10.1073/pnas.1915255117. PubMed PMID: 32198200; PMCID: PMC7149309.

61. Pothoulakis C, Gilbert RJ, Cladaras C, Castagliuolo I, Semenza G, Hitti Y, Montcrief JS, Linevsky J, Kelly CP, Nikulasson S, Desai HP, Wilkins TD, LaMont JT. Rabbit sucrase-isomaltase contains a functional intestinal receptor for Clostridium difficile toxin A. J Clin Invest. 1996;98(3):641–9. Epub 1996/08/01. doi: 10.1172/JCI118835. PubMed PMID: 8698855; PMCID: PMC507473.

62. Na X, Kim H, Moyer MP, Pothoulakis C, LaMont JT. gp96 is a human colonocyte plasma membrane binding protein for Clostridium difficile toxin A. Infect Immun. 2008;76(7):2862–71. Epub 2008/04/16. doi: 10.1128/IAI.00326-08. PubMed PMID: 18411291; PMCID: PMC2446715.

63. Tao L, Tian S, Zhang J, Liu Z, Robinson-McCarthy L, Miyashita SI, Breault DT, Gerhard R, Oottamasathien S, Whelan SPJ, Dong M. Sulfated glycosaminoglycans and low-density lipoprotein receptor contribute to Clostridium difficile toxin A entry into cells. Nat Microbiol. 2019. Epub 2019/06/05. doi: 10.1038/s41564-019-0464-z. PubMed PMID: 31160825.

64. Schöttelndreier D, Langejürgen A, Lindner R, Genth H. Low Density Lipoprotein Receptor-Related Protein-1 (LRP1) Is Involved in the Uptake of Clostridioides difficile Toxin A and Serves as an Internalizing Receptor. Front Cell Infect Microbiol. 2020;10:565465. Epub 20201019. doi: 10.3389/fcimb.2020.565465. PubMed PMID: 33194803; PMCID: PMC7604483.

65. LaFrance ME, Farrow MA, Chandrasekaran R, Sheng J, Rubin DH, Lacy DB. Identification of an epithelial cell receptor responsible for Clostridium difficile TcdB-induced cytotoxicity. Proc Natl Acad Sci U S A. 2015;112(22):7073–8. Epub 20150518. doi: 10.1073/pnas.1500791112. PubMed PMID: 26038560; PMCID: PMC4460460.

66. Tao L, Zhang J, Meraner P, Tovaglieri A, Wu X, Gerhard R, Zhang X, Stallcup WB, Miao J, He X, Hurdle JG, Breault DT, Brass AL, Dong M. Frizzled proteins are colonic epithelial receptors for C. difficile toxin B. Nature. 2016;538(7625):350-5. Epub 2016/10/21. doi: 10.1038/nature19799. PubMed PMID: 27680706; PMCID: PMC5519134.

67. Yuan P, Zhang H, Cai C, Zhu S, Zhou Y, Yang X, He R, Li C, Guo S, Li S, Huang T, Perez-Cordon G, Feng H, Wei W. Chondroitin sulfate proteoglycan 4 functions as the cellular receptor for Clostridium difficile toxin B. Cell Res. 2015;25(2):157–68. Epub 2014/12/31. doi: 10.1038/cr.2014.169. PubMed PMID: 25547119; PMCID: PMC4650570.

68. Diaz-Gonzalez F, Milano M, Olguin-Araneda V, Pizarro-Cerda J, Castro-Cordova P, Tzeng SC, Maier CS, Sarker MR, Paredes-Sabja D. Protein composition of the outermost exosporium-like layer of *Clostridium difficile* 630 spores. J Proteomics. 2015;123:1–13. Epub 2015/04/08. doi: 10.1016/j.jprot.2015.03.035. PubMed PMID: 25849250.

69. Barketi-Klai A, Hoys S, Lambert-Bordes S, Collignon A, Kansau I. Role of fibronectin- binding protein A in *Clostridium difficile* intestinal colonization. J Med Microbiol. 2011;60(Pt 8):1155–61. Epub 2011/02/26. doi: 10.1099/jmm.0.029553-0. PubMed PMID: 21349990.

70. Lin YP, Kuo CJ, Koleci X, McDonough SP, Chang YF. Manganese binds to Clostridium difficile Fbp68 and is essential for fibronectin binding. J Biol Chem. 2011;286(5):3957–69. Epub 20101109. doi: 10.1074/jbc.M110.184523. PubMed PMID: 21062746; PMCID: PMC3030396.

71. Peritore-Galve FC, Shupe JA, Cave RJ, Childress KO, Washington MK, Kuehne SA, Lacy DB. Glucosyltransferase-dependent and independent effects of Clostridioides difficile toxins during infection. PLoS Pathog. 2022;18(2):e1010323. Epub 20220217. doi: 10.1371/journal.ppat.1010323. PubMed PMID: 35176123; PMCID: PMC8890742.

72. Kuehne SA, Collery MM, Kelly ML, Cartman ST, Cockayne A, Minton NP. Importance of toxin A, toxin B, and CDT in virulence of an epidemic Clostridium difficile strain. J Infect Dis. 2014;209(1):83–6. Epub 20130809. doi: 10.1093/infdis/jit426. PubMed PMID: 23935202; PMCID: PMC3864386.

73. Kuehne SA, Cartman ST, Minton NP. Both, toxin A and toxin B, are important in *Clostridium difficile* infection. Gut Microbes. 2011;2(4):252–5. Epub 2011/07/01. doi: 10.4161/gmic.2.4.16109. PubMed PMID: 21804353; PMCID: PMC3260544.

74. Mohamed MFH, Ward C, Beran A, Abdallah MA, Asemota J, Kelly CR. Efficacy, Safety, and Cost-effectiveness of Bezlotoxumab in Preventing Recurrent Clostridioides difficile Infection : Systematic Review and Meta-analysis. J Clin Gastroenterol. 2024;58(4):389–401. Epub 20240401. doi: 10.1097/mcg.0000000000001875. PubMed PMID: 37395627.

75. Johnson TM, Molina KC, Howard AH, Schwarz K, Allen L, Huang M, Bajrovic V, Miller MA. Real-World Comparison of Bezlotoxumab to Standard of Care Therapy for Prevention of Recurrent Clostridioides difficile Infection in Patients at High Risk for Recurrence. Clin Infect Dis. 2021. Epub 20210804. doi: 10.1093/cid/ciab674. PubMed PMID: 34665248.

76. Yang Z, Ramsey J, Hamza T, Zhang Y, Li S, Yfantis HG, Lee D, Hernandez LD, Seghezzi W, Furneisen JM, Davis NM, Therien AG, Feng H. Mechanisms of protection against Clostridium difficile infection by the monoclonal antitoxin antibodies actoxumab and bezlotoxumab. Infect Immun. 2015;83(2):822–31. Epub 20141208. doi: 10.1128/iai.02897-14. PubMed PMID: 25486992; PMCID: PMC4294251.

77. Warn P, Thommes P, Sattar A, Corbett D, Flattery A, Zhang Z, Black T, Hernandez LD, Therien AG. Disease Progression and Resolution in Rodent Models of Clostridium difficile Infection and Impact of Antitoxin Antibodies and Vancomycin. Antimicrob Agents Chemother. 2016;60(11):6471–82. Epub 20161021. doi: 10.1128/aac.00974-16. PubMed PMID: 27527088; PMCID: PMC5075051.

78. Chandrasekaran R, Kenworthy AK, Lacy DB. *Clostridium difficile* Toxin A Undergoes Clathrin-Independent, PACSIN2-Dependent Endocytosis. PLoS Pathog. 2016;12(12):e1006070. Epub 2016/12/12. doi: 10.1371/journal.ppat.1006070. PubMed PMID: 27942025; PMCID: PMC5152916.

79. Castro-Córdova P, Díaz-Yáñez F, Muñoz-Miralles J, Gil F, Paredes-Sabja D. Effect of antibiotic to induce *Clostridioides difficile*-susceptibility and infectious strain in a mouse model of *Clostridioides difficile* infection and recurrence. Anaerobe. 2020;62:102149. Epub 2020/01/12. doi: 10.1016/j.anaerobe.2020.102149. PubMed PMID: 31940467.

80. Pizarro-Guajardo M, Diaz-Gonzalez F, Alvarez-Lobos M, Paredes-Sabja D. Characterization of Chicken IgY Specific to *Clostridium difficile* R20291 Spores and the Effect of Oral Administration in Mouse Models of Initiation and Recurrent Disease. Front Cell Infect Microbiol. 2017;7:365. Epub 2017/09/01. doi: 10.3389/fcimb.2017.00365. PubMed PMID: 28856119; PMCID: PMC5557795.

81. Castro-Córdova P, Mendoza-León MJ, Paredes-Sabja D. Using a ligate intestinal loop mouse model to investigate Clostridioides difficile adherence to the intestinal mucosa in aged mice. PLoS One. 2021;16(12):e0261081. Epub 20211222. doi: 10.1371/journal.pone.0261081. PubMed PMID: 34936648; PMCID: PMC8694449.

82. Calderon-Romero P, Castro-Cordova P, Reyes-Ramirez R, Milano-Cespedes M, Guerrero- Araya E, Pizarro-Guajardo M, Olguin-Araneda V, Gil F, Paredes-Sabja D. *Clostridium difficile* exosporium cysteine-rich proteins are essential for the morphogenesis of the exosporium layer, spore resistance, and affect *C. difficile* pathogenesis. PLoS Pathog. 2018;14(8):e1007199. Epub 2018/08/09. doi: 10.1371/journal.ppat.1007199. PubMed PMID: 30089172.

83. Johansson ME, Larsson JM, Hansson GC. The two mucus layers of colon are organized by the MUC2 mucin, whereas the outer layer is a legislator of host-microbial interactions. Proc Natl Acad Sci U S A. 2011;108 Suppl 1(Suppl 1):4659-65. Epub 20100625. doi: 10.1073/pnas.1006451107. PubMed PMID: 20615996; PMCID: PMC3063600.

